# Parallel Evolution of Salinity Tolerance in *Arabidopsis thaliana* Accessions from Cape Verde Islands

**DOI:** 10.1101/2024.05.21.595092

**Authors:** Félix J Martínez Rivas, Dorothee Wozny, Zeyun Xue, Elodie Gilbault, Thomas Sapir, Melissa Rouille, Antony Ricou, Joaquín Medina, Laurent D. Noël, Emmanuelle Lauber, Aline Voxeur, Olivier Loudet, Gilles Clément, Jose M Jiménez-Gómez

## Abstract

Soil salinization poses a significant threat to crop production impacting one fifth of all cultivated land. The Cape Verde Islands are located 600 km from the coast of Africa and are characterized by high salinity soils and inland water sources.

In this study we find that *Arabidopsis thaliana* plants native to these islands accumulate a metabolite that protects them from salt stress. We partially characterized this metabolite as glucuronyl-mannose. We find that the ability to produce glucuronyl-mannose evolved independently in two different islands from the same archipelago through mutations in the same gene, an alpha glycosidase protein that we named GH38cv. These cases of parallel evolution suggest positive selection towards the increase of salt tolerance with low fitness costs. Indeed, plants carrying derived alleles of GH38cv do not present growth defects or low defenses under normal conditions, but show better germination rates, longer roots and better hydric status than non-mutated plants when exposed to salt stress. These findings provide a knowledge-based method to develop salt resilient crops using natural mechanisms, which could be attractive both to conventional and organic agriculture.

## Introduction

Plant species colonizing harsh ecological niches can evolve characteristics that help them adapt to local challenges. Parallel evolution occurs when two lineages independently evolve these characteristics and is considered a signal of positive selection with low fitness costs (Woods et al., 2006; Kreiner et al., 2019; Konečná et al., 2021). For these reasons, identifying these mechanisms for traits of agricultural interest provides valuable knowledge to increase crop tolerance to environmental threats under a changing climate.

Soil salinization is spreading globally and affects 25% of irrigated lands (Shahid et al., 2018). Only 1% of all plants can tolerate high salt concentrations, complicating cultivation in saline areas such as islands or coastal regions (Garcia-Caparros et al., 2023). The Cape Verde Islands are located 600 km from its closest mainland off the coast of Africa and have high salinity both in their soil and inland water sources (Zhao and Filker, 2018; Cruz et al., 2023). The Cvi-0 accession of the model plant *Arabidopsis thaliana* was collected in these islands in the 1990s and has been extensively studied because of its unique characteristics (Alonso-Blanco et al., 2003; Alonso-Blanco et al., 2005; Jakobson et al., 2016; Durand et al., 2021; Shahzad et al., 2024). More than 20 years later, another 335 individuals from seven populations in the islands of Santo Antão and Fogo were collected (Fulgione et al., 2022).

These individuals were more closely related to one another than to any accession in the worldwide collection. Their ancestors were predicted to colonize the island of Santo Antão between five and seven thousand years ago, and from there, the island of Fogo between three and five thousand years ago (Fulgione et al., 2022). During this time, plants in these islands have diverged in traits of high adaptive value such as flowering time, nutrient transport and water use efficiency (Fulgione et al., 2022; Tergemina et al., 2022; Elfarargi et al., 2023; Neto and Hancock, 2023).

Plant responses to high salinity have been well characterized at the physiological (van Zelm et al., 2020) and molecular levels (Zhou et al., 2024). Accumulation of toxic ions impairs nutrient uptake, reduces cell turgor, induces stomatal closure and alters photosynthesis (Zhou et al., 2024). Reactive oxygen species production and oxidative stress in the cells trigger drastic metabolic changes directed to reduce water loss and enhance cell turgor (Yang and Guo, 2018). Using techniques such as liquid chromatography-mass spectrometry or gas chromatography-mass spectrometry, we can accurately quantify hundreds of metabolites involved in these processes (Alseekh et al., 2021). When metabolomics is performed in large plant populations, the loci involved in the differential accumulation of metabolites can be mapped to chromosomal regions (Lisec et al., 2008; Brotman et al., 2011; Joseph et al., 2014; Wu et al., 2016; Knoch et al., 2017; Wu et al., 2018; Naake et al., 2024). Once the precise mutations involved in production of metabolites are identified, gene editing techniques can be harness to reproduce these mutations in plants of societal interest.

In this work we conducted metabolic profiling in a population derived from a cross between the reference accession Col-0 and the Cape Verde Islands accession Cvi-0. We found a disaccharide that protects Arabidopsis plants from salt stress, and we mapped the mutation that led to the generation of this metabolite to the alpha-mannosidase gene AT3G26720, which we named GH38cv. We analyzed the genome of additional plants from other islands and we identified a case of parallel evolution in GH38cv, suggesting positive selection for this tolerance mechanism in the archipelago.

## Results

### Metabolomics in the Col-0 x Cvi-0 RIL population

We explored metabolite variation in a recombinant inbred line (RIL) population derived from the refence *Arabidopsis thaliana* accession Col-0, from central Europe, and Cvi-0, an accession from Cape Verde Islands (Simon et al., 2008) (Suppl dataset 1). To ensure robustness of the metabolic dataset, two replicates of each line were grown in two consecutive and independent experiments in the Phenoscope, a robotic platform that automatically waters and monitors plants several times a day (Tisné et al., 2013). The aerial part of each plant was collected after 23 days on the robot (31 days after sowing), and we used GC-MS for metabolite quantification. We identified 118 metabolites in more than half of the lines in the population (Suppl dataset 1). Clustering of the metabolic profiles across the population showed higher correlations between primary metabolites, and between metabolites in the TCA cycle (Figure S1).

We used mixed effect models to estimate the abundance of each metabolite in each line in the RIL population removing the effect of the batch of injection into the GC-MS machine (see methods). Corrected values were used to perform quantitative trait locus (QTL) analysis. We detected 44 QTLs for 35 metabolites, an average of 1.25 QTLs per metabolite with a detectable genetic basis (Figure S2, Figure S3A, Suppl dataset 2). We detected QTL hotspots in chromosome 1 and chromosome 3 (Figure S3B), and Cvi-0 alleles at both hotspots increased concentration of metabolites such as aconitate, citrate, maleate, phosphate and myo-inositol-1-P (Figures S2 and S3). Among the top ten highest effect QTLs (Figure 1A), the QTL for proline in chromosome 2 and the QTL affecting a glucosinolate derivate in chromosome 3 had been previously described (Zhang et al., 2006; Kesari et al., 2012). The two strongest QTLs were both located in chromosome 3 and affected glucuronic acid and another metabolite with a retention index of 2745.9 and a characteristic ion with a m/z of 494. We called this metabolite U2746 based on its retention index. In the RIL population, Cvi-0 alleles at the U2746 QTL in chromosome 3 decreased the abundance of glucuronic acid and increased the abundance of U2746, suggesting that a single causal mutation controls both metabolites.

**Figure 1.**
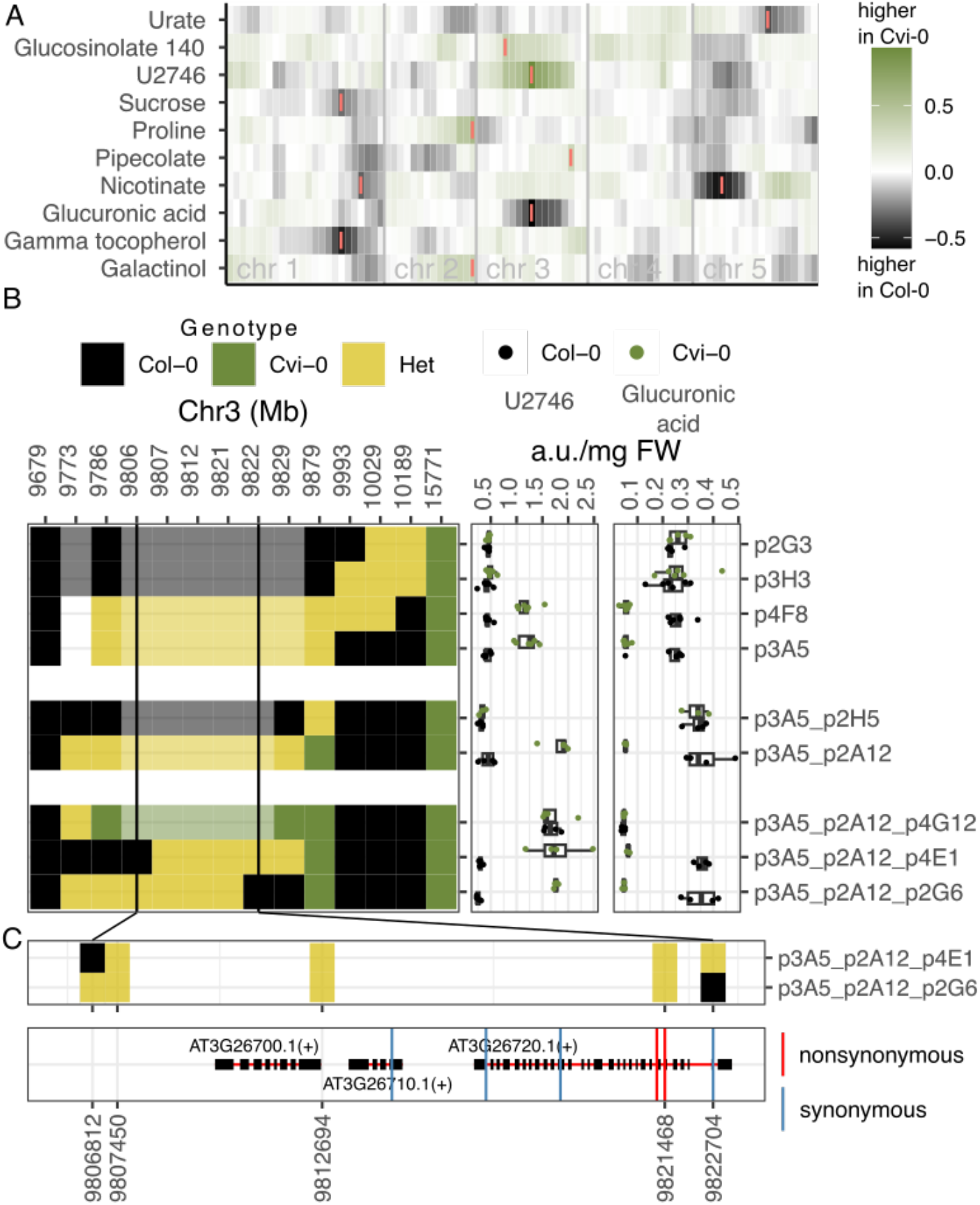
Mapping and fine mapping of metabolite QTLs in Cvi-0 x Col-0. (A) Heatmap with QTL effects for the top ten metabolite QTLs in the population. Markers are placed along the x axis, separated by chromosomes by vertical gray bars. The color scale in the heatmap represent the calculated additive effects of the homozygous alleles. The marker with the highest LOD score for each trait is marked with colored vertical bars. (B) Left panel shows the genotypes of recombinant plants selected from HIF474 (y axis) for markers at the indicated positions in chromosome 3 (x axis). Solid and light coloring represents real and imputed genotypes respectively. Vertical black lines indicate the region delimiting the candidate mutation. Right panel shows the abundance of U2746 and glucuronic acid in the progeny of each recombinant plants shown in the left. (C) Top panel shows the genotype and physical position of the markers tested in the two recombinants that delimit the causal region. Bottom panel shows the genes in the causal region and the position of all coding mutations between Col-0 and Cvi-0.

### Fine mapping of the U2746 QTL

To fine map the mutation causing the accumulation of U2746 in Cvi-0 we grew heterogenous inbred families (HIFs) that contain a single genomic fragment in heterozygosis overlapping the region of the QTL (Loudet et al., 2005). The progeny of two among the three HIFs selected showed differences in the accumulation of U2746 associated to contrasting alleles of the QTL (Figure S4). We screened close to 1500 descendants from one of these HIFs (8HV474) during three rounds of fine mapping, which led us to delimit the QTL region down to three genes (Figure 1B). During all fine mapping experiments, metabolic profiling of U2746 and glucuronic acid in lines carrying homozygous Col-0 and Cvi-0 alleles at the QTL region showed remarkable negative correlation (Figure 1B). For validation of the QTL we selected the homozygous descendants from HIF line p3A5_p2A12_p4E1, which we hereby name HIF-Col and HIF-Cvi. One of the three candidate genes considered encodes for a glycoside hydrolase family 38 protein (AT3G26720, hereafter named GH38cv). This protein has been described to function as an alpha-mannosidase (Fujiyama et al., 2001), and has been shown to increase salt resistance both in Arabidopsis and wheat (Wang et al., 2020b; Wang et al., 2020a). We thus investigated the implication of GH38cv in the differential accumulation of U2746 and glucuronic acid in Col-0 and Cvi-0.

### GH38cv underlies the U2746 QTL

We first confirmed the causal link between GH38cv and U2746 accumulation by using two independent T-DNA lines with insertions in the *GH38cv* gene. Both lines showed increased levels of U2746 and decreased levels of glucuronic acid with respect to their wild type Col-0 (Figure 2A and 2B). Remarkably, one of the mutants had an insertion in an intron and showed lower levels of U2746 than the mutant with an insertion in an exon, which agrees with the latter being more disruptive of the activity of the gene since it changes its amino acid sequence (Wang, 2008).

**Figure 2.**
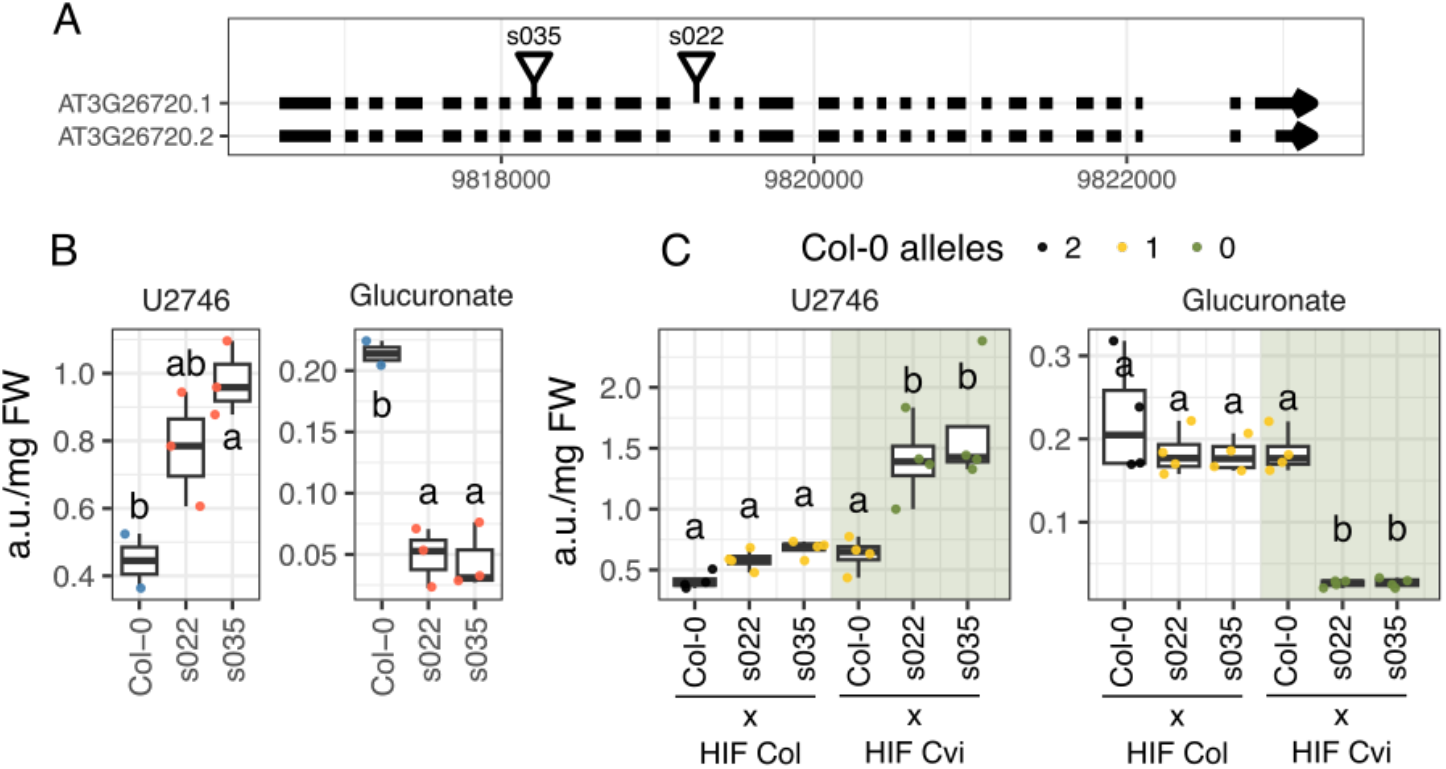
Confirmation of the effect of AT3G26720 on metabolite accumulation. (A) Genomic positions of T-DNA insertions in AT3G26720 based on coordinates from Salk Signal T-DNA Express in the TAIR10 genome reference and annotation. Black boxes are exons, arrows point to the direction of transcription. s035 stands for line SALK_035007C and s022 for SALK_022207C. (B) Normalized abundance in arbitrary units of U2746 (left) and glucuronic acid (right) for Col-0 and two T-DNA insertion mutants. (C) Normalized abundance in arbitrary units of U2746 (left) and glucuronic acid (right) for hybrid plants resulting from crosses of the specified lines in the x axis. Different letters represent significant differences between lines (one way ANOVA, Tukey HSD test, p<0.05). Shaded area in each plot indicate lines that carry one Cvi-0 allele and non-shaded areas indicate lines with no Cvi-0 alleles. Dot color mark the number of Col-0 alleles in each line.

We then used quantitative complementation to confirm the differential effect of the Col-0 and the Cvi-0 alleles of GH38cv on the accumulation of U2746 and glucuronic acid. For this, we crossed the T-DNA insertion mutants with the homozygous HIF-Col and HIF-Cvi lines to obtain plants displaying 0, 1 or 2 copies of the Col-0 allele, and 0 or 1 copies of the Cvi-0 allele. Variation in the number of GH38cv Col-0 alleles have an effect in U2746 and glucuronic acid accumulation, while the number of Cvi-0 alleles had no effect (Figure 2C). This suggests that the Cvi-0 alleles of GH38cv are null. In summary, these experiments indicate that GH38cv directly controls the accumulation of U2746 and glucuronic acid, and that the Cvi-0 allele, but not the Col-0 allele, carries one or several mutations that compromise its function.

### Identification of U2746 as glucuronyl-mannose

We then studied the nature of the U2746 metabolite and its relation with glucuronic acid. Analysis of the GC-MS data acquired from the RIL population revealed similarities in the retention index and mass spectra between U2746 and the disaccharide maltose (Figure S5). Their fragmentation spectra differed in the shift of a peak in U2746 from m/z 480 to m/z 494, representing a 14 Da difference that could be explained by the presence of uronic acid as one of the saccharides (Figure 3). To confirm the disaccharide nature of U2746, we performed acid hydrolysis of ether bonds in HIF-Col and HIF-Cvi extracts (Fig 3B).

**Figure 3.**
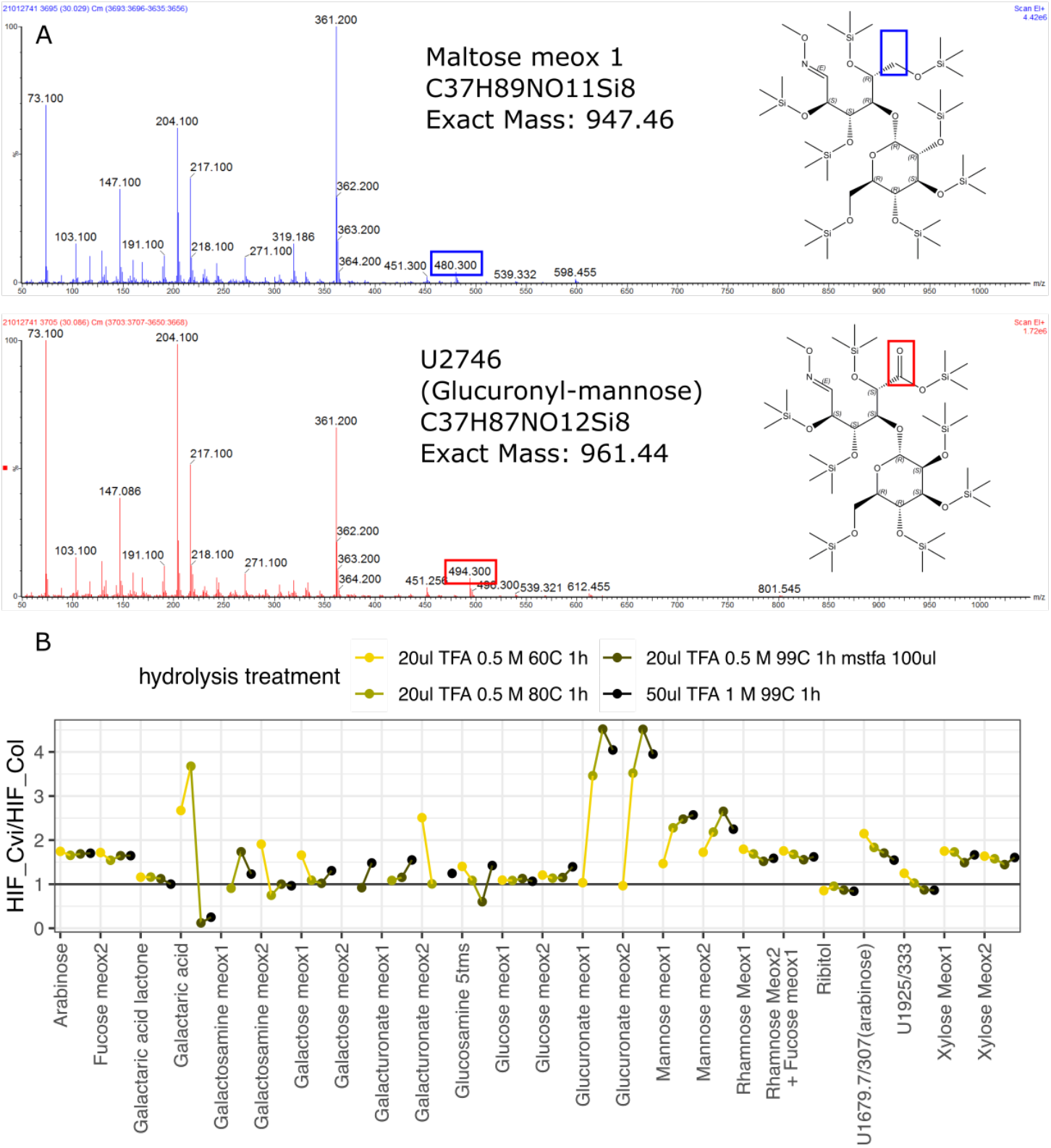
Characterization of U2746. (A) EI 70ev spectra for maltose (top) and U2746 (bottom) from m/z 50 to 1050 from leaf extracts of HIF-Cvi. The main difference in these spectra is marked with a box. This 14 Da difference fits with U2746 having an extra oxygen, marked in the representation of the molecules to the right of the spectra (B) Ratio of sugars released after acid hydrolysis of leaf extracts from HIF-Col and HIF-Cvi. Glucuronate and mannose are released in higher concentrations in HIF-Cvi than in HIF-Col upon TFA treatment, suggesting that U2746 is a glucuronyl-mannose.

Disappearance of U2746 lead to increase abundance of methoxyamine derivatives of glucuronic acid and mannose, suggesting mannose as the second saccharide.

We then purified leaf extracts from Col-0 and one of the T-DNA insertion lines with an anion-exchange column. U2746 eluted with all the organic acids, proving that it contains an acidic moiety (Figure S6A). Upon TFA acid hydrolysis of the partially purified sample, the disappearance of U2746 molecule led to the accumulation of neutral and acidic monosaccharides (hexoses in Figure S6B). Among the released sugars, mannose and glucuronic acid were more abundant in the T-DNA insertion line than in Col-0, supporting again that U2746 is glucuronyl-mannose (Figure S6C). During all QTL and fine mapping experiments we found a tight correlation between the concentration of U2746 and glucuronic acid, but we did not find correlations between U2746 and mannose or any other saccharide (Figure S7). One possibility is that mannose is metabolized faster than glucuronic acid, for example towards ascorbic acid synthesis (Conklin et al., 1999). Another possibility is that mannose is replaced by another hexose that could not be detected in the eluate of the exchange columns. In summary, we propose that U2746 is an ether linked glucuronyl-mannose, and its overaccumulation in Arabidopsis depends on the presence of an inactive GH38cv protein.

### Characterizing the role of GH38cv and U2746 in Arabidopsis

We explored the dynamics in the abundance of both U2746 and glucuronic acid during plant development. The concentration of U2746 increased with plant age up to four times more in HIF-Cvi than in HIF-Col, and is negatively correlated with the concentration glucuronic acid until senescence, when HIF-Cvi lines accumulate both glucuronic acid and U2746 (Figure S8). The HIFs do not show differences in mannose during their lifespan (Figure S8).

It has been reported that mutations in GH38cv increase germination rate, root length and fresh weight in Arabidopsis under high salinity (Wang et al., 2020a; Wang et al., 2020b). We confirmed that the natural Cvi-0 mutation had a similar effect using the HIF lines segregating for the mutated GH38cv allele. In our conditions, HIF-Cvi had better germination rates, longer roots, and higher stomatal conductance than HIF-Col when exposed to salt stress (Figure 4 and S12).

**Figure 4.**
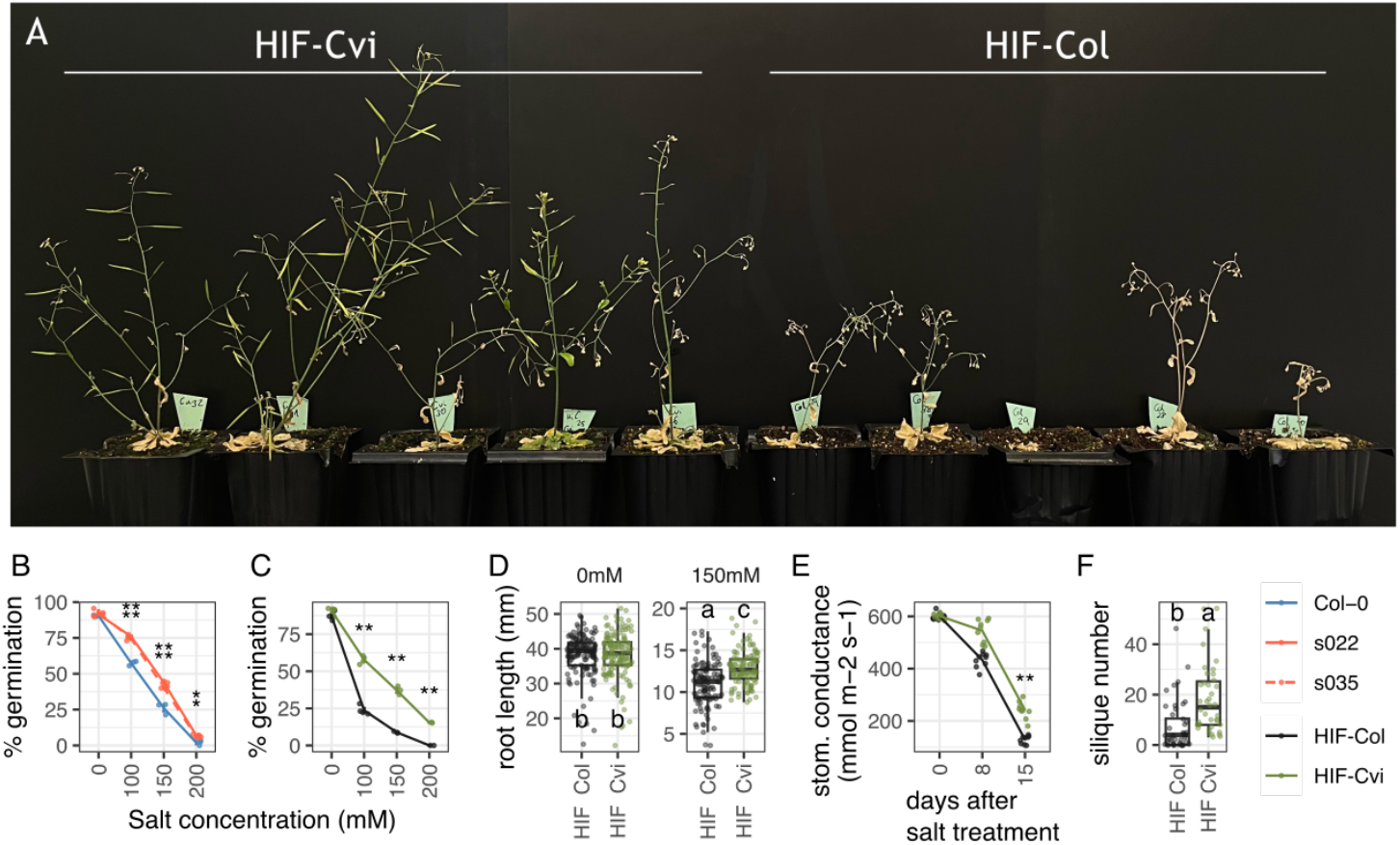
Effect of mutations in AT3G26720 in responses to high salinity. (A) HIF-Col and HIF-Cvi plants after 7 days of irrigation with salted water. (B) and (C) Percent germination after 7 days in different concentrations of salt in seeds from T-DNA insertion lines (B) or HIF lines with contrasting alleles of AT3G26720. Asterisks indicate significant differences at a given salt concentration between Col-0 and the T-DNA insertion mutants in (B) or Col-0 and Cvi-0 alleles in the HIFs in (C) (Student t-test ** p<0.01, * p<0.05). Data are from 2 independent experiments. (D) Root length 5 days after transfer to control (0mM) or salt medium (150mM) in the HIF lines with contrasting GH38cv alleles. Data are from 4 independent experiments. Different letters indicate significant differences between lines and conditions (two-way ANOVA, Tukey HSD test, p<0.05). (E) Stomatal conductance in plants treated with salted water. Treatment starts at day 0. Asterisks indicate significant differences at a given day (Student t-test ** p<0.01). (F) Silique number in plants from (A) (p=7.15e-5).

**Figure 6.**
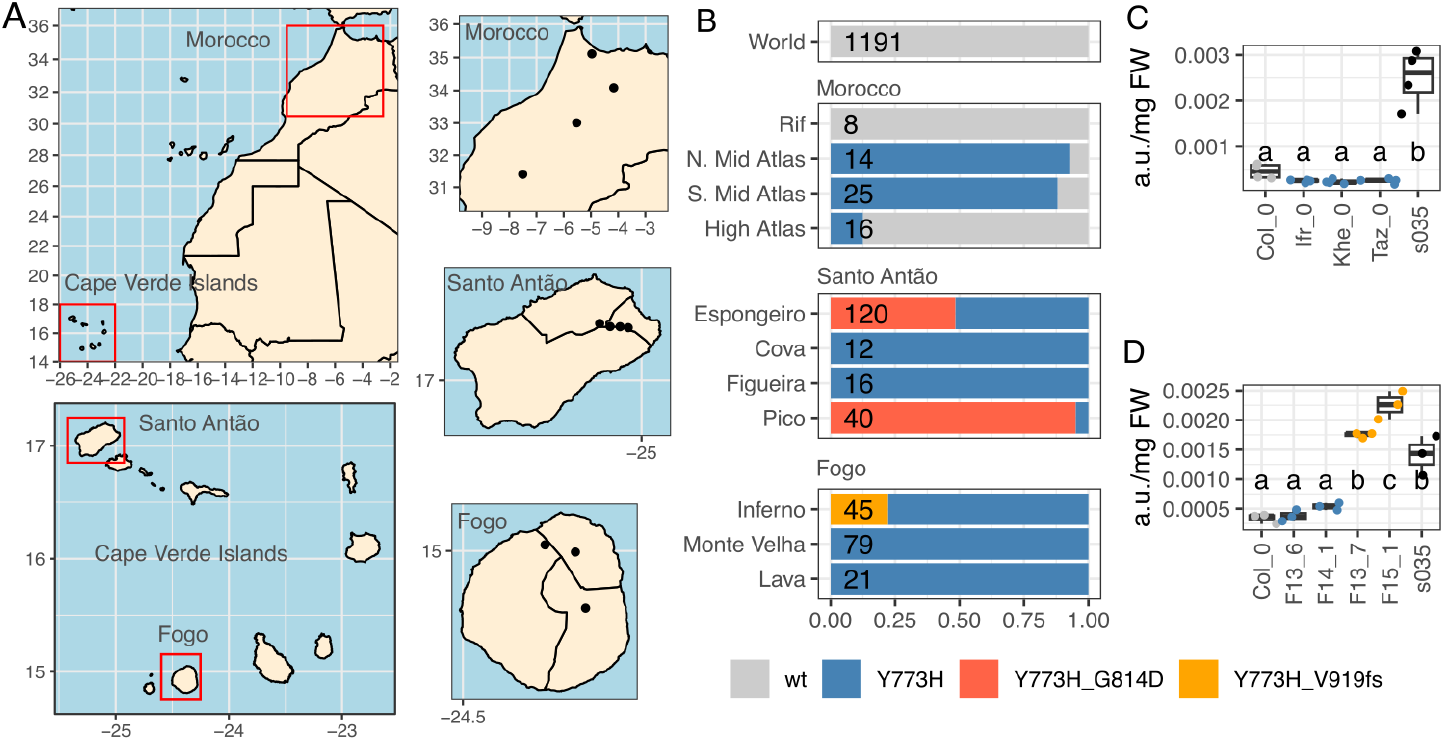
Distribution in Cape Verde Islands and Morocco of variants of interest in GH38cv. (A) Maps with geographic regions of interest. Dots indicate the places the populations were collected. (B) Frequency of variants Y773H, G814D and V919fs found for GH38cv in the populations shown in Morocco, Santo Antão, and Fogo. Plants with Col-0 alleles for these three variants are labelled wt. The number of individuals in each population is shown in each bar. (C and D) U2746 abundance in arbitrary units for accessions from Morocco (C) or the island of Fogo (D). Different letters indicate significant differences between lines (one-way ANOVA, Tukey HSD test, p<0.05).

We also studied whether the accumulation of the U2746 metabolite compromised other aspects of plant growth or stress responses, which would prevent its practical application in agriculture. We did not observe any visual differences between HIF-Col and HIF-Cvi, or between Col-0 and the T-DNA mutants, when grown under standard conditions. We also analyzed the images taken daily from the Cvi-0 x Col-0 RILs in the Phenoscope robot during the QTL analysis experiments. We first confirmed that RILs with contrasting alleles at GH38cv, presented significant differences in the accumulation of the U2746 metabolite and glucuronic acid, but not mannose (Figure S9). RILs carrying Cvi-0 alleles at GH38cv, thus accumulating U2746, showed small but significantly larger rosette area, increased growth rates, and their leaves had higher intensity of red and green colours (Figure S10). We then tested whether accumulation of U2746 affected biotic interactions with virulent bacteria (*Xanthomonas campestris* pv. *campestris*) or fungi (*Botrytis cinerea*). HIF-Cvi plants did not show significant differences with HIF-Col plants in their resistance to pathogens (Figure S11). These experiments allow us to conclude that the differential accumulation of the U2746 metabolite due to mutations in GH38cv confers plants with an advantage under high salt concentrations, but does not compromise overall plant health under standard conditions.

### Genetics of GH38cv in Arabidopsis populations

We then studied the origin of the GH38cv allele from the Cape Verde Islands. Cvi-0 carries two nonsynonymous mutations in GH38cv with respect to the Col-0 reference: a tyrosine to histidine change at position 773 of the protein (Y773H), and a glycine to aspartic acid change at position 814 (G814D) (Figure 1). Both positions are invariable in orthologous genes from 11 *Brasicaceae* species, but position G814D is more conserved than Y773D across a diverse set of plant families (Figure S13). We explored the distribution of these mutations in short read data from 1,607 worldwide Arabidopsis accessions accessible in public repositories (1001 Genomes Consortium, 2016), including regional sets from the Cape Verde Islands (Fulgione et al., 2022) and nearby regions such as Northeast Africa (Durvasula et al., 2017) or the Iberian peninsula (Arteaga et al., 2021). The Y773H and G814D mutations are only present in accessions from Africa (Figure 5, Suppl dataset 3). The Y773H substitution is fixed in all accessions from Cape Verde Islands and is found in three out of four Moroccan populations, implying that it was likely present in the accessions that first colonized the Cape Verde Islands (Figure 5B). Accessions from Morocco that carry the Y773H mutation do not accumulate U2746, indicating that the G184D mutation is the most plausible candidate to affect the function of GH38cv (Figure 5C). The G814D mutation appears in two out of four populations collected in the island of Santo Antão, suggesting it originated in this island. It is present in 79% of the individuals sampled at Pico da Cruz, and in 48% of the individuals sampled in Espongeiro, where we also found one plant carrying the mutation in heterozygous state (Figure 5B).

We then looked for other alleles of GH38cv that could indicate selection of this gene during adaptation to other regions worldwide. We used the short-read sequence data to reconstruct 92 different alleles of GH38cv (Figure S14). We only found one non-functional allele, which contained a 1 bp deletion producing a shift in its open reading frame (V919fs mutation, allele a21 in Figure S14). Interestingly, this allele also originates from the Cape Verde Islands, albeit from the island of Fogo instead of the island of Santo Antão, where the Cvi-0 allele originates (Suppl dataset 3). The frameshift allele was present in ten out of the forty-five individuals from the Inferno population of Fogo (Figure5B). Individuals from Fogo carrying the V919fs mutation showed significantly higher concentrations of U2746 than neighboring plants without the mutation (Figure 5D).

In summary, we found two nonfunctional mutations that arose independently in the Cape Verde Islands, targeting the same gene and promoting the accumulation of U2746, thus increasing their tolerance to high salinity. The presence of these two independent mutational events in the same geographic region suggests positive selection on GH38cv to induce accumulation of this metabolite and increase plant tolerance to the high salinity conditions in the islands.

## Discussion

High salinity in soil and irrigation water negatively affects plant growth and crop productivity. We investigated natural strategies adopted by plants to colonize the high salinity soils on the Cape Verde Islands, and we found that accumulation of U2746 improves salt tolerance. This metabolite is present at low concentration in most Arabidopsis accessions, but significantly accumulates in GH38cv loss of function mutants. Although GH38 remains largely uncharacterized, numerous proteomics studies report that it is secreted to the cell wall where it is likely active, as summarized in the SUBA5 database (Hooper et al., 2017). As GH38 is predicted to exhibit alpha-mannosidase activity, it could be expected that GH38 targets mannose-rich carbohydrates, such as hemicelluloses, for restructuring or degradation. The loss of GH38 activity causes an accumulation of glucoronyl-mannose and a decrease in glucuronic acid concentration, while mannose concentration remains unaffected. Glucuronic acid is an important component of xylan, the main structural hemicelluloses in plant cell walls (Oliveira, 2023). Xylan content has been shown to decrease in the cell wall upon salt stress treatment (Oliveira et al., 2020). It is possible that accumulation of glucuronyl mannose in Cape Verde Island accessions limits the availability of glucuronic acid for proper cell wall modelling, thus affecting salt tolerance.

Functional studies to characterize GH38cv activity and substrate specificity, as well as mutant cell wall composition analysis, will aid in revealing the mechanism driving the altered carbohydrate profile in gh38cv mutants.

Another potential mechanism for salinity tolerance may stem from glucuronyl mannose acting as an osmoprotectant. Other disaccharides such as trehalose have well known roles in salinity stress (Nawaz et al., 2022). Trehalose has been detected at high concentrations in halophyte plants, where it seems to act as an osmolyte (Bianchi et al., 1993), increasing water uptake potential and protecting cellular membranes against oxidative damage (Balasubramaniam et al., 2023). However, in non-halophyte plants trehalose is found at very low quantities and is thought to have a role as a signaling molecule, although no direct proof exists (Fernandez et al., 2010; Yang et al., 2022). In any case, trehalose application or endogenous production has proven effective to enhance salinity tolerance in many economically important species, such as wheat, rice, maize, or tomato (Abdallah et al., 2016; Rohman et al., 2019; Sadak, 2019; Yang et al., 2022). Further research is needed to conclude whether accumulation of glucuronyl-mannose has a role as an osmolyte, a cell wall modifier, a signaling molecule, or all of the above.

In this study we discover that the accumulation of glucuronyl-mannose has independently evolved twice in Arabidopsis plants from the Cape Verde Islands. Their location far from the coast, their volcanic soil and harsh weather, and their relatively early colonization by Arabidopsis provide an outstanding opportunity to study adaptation to challenging environments. Previous works with populations from these islands have revealed cases of parallel evolution towards early flowering and adapted nutrient transport (Fulgione et al., 2022; Tergemina et al., 2022). In both cases, independent mutations in one of two flowering time genes (FRI or FLC) or nutrient transport genes (IRT1 and NRAMP1) contributed to the evolution of new phenotypes in the islands. We find a similar case of parallel evolution, although selection occurred in a single gene (GH38cv). Examples of genomic regions with high frequencies of adaptive mutations have been reported in sticklebacks, plants and other organisms (Cockram et al., 2007; Chan et al., 2010; Xie et al., 2019). In Arabidopsis, allelic heterogeneity has also been reported among local populations (Barboza et al., 2013; Saez-Aguayo et al., 2014). At the evolutionary level, the reiterative mutation of GH38cv can be taken as evidence of positive selection and low fitness costs of the mutated alleles in the newly colonized environment (Chan et al., 2010).

Our findings can have practical implications for advances in agriculture. Disaccharides such as trehalose can been exogenously applied to crops to alleviate the negative effects of soil salinity (Abdallah et al., 2016; Rohman et al., 2019; Sadak, 2019; Yang et al., 2022). With further testing, U2746 may reveal similar potential as a naturally derived soil conditioner. Additionally, it would be possible to edit *GH38cv* homologs to confer salinity tolerance in globally important crop varieties, as *GH38cv* homologs are present in a wide number of plant families (Figure S13). In wheat, GH38cv interacts with the U-box E3 ubiquitin ligase TaPUB1, and its expression is induced by salt treatment (Wang et al., 2020a). *GH38cv* overexpression in the grass *Brachypodium distachyon* resulted in decreased tolerance to salinity, confirming its role in salt stress responses (Wang et al., 2020b; Wang et al., 2020a). In summary, both the synthesis and application of glucuronyl-mannose, as well as the inactivation of GH38cv through gene editing in crops open new avenues to improve salinity tolerance in crops and ornamental plants using technology-based methods inspired by natural processes.

## Methods

### Plant growth for QTL analysis and confirmation

A subset of 90 lines from a Cvi-0 × Col-0 recombinant inbred line (RIL) population (Simon et al., 2008) was used for metabolite profiling and mapping of metabolomic quantitative trait loci (QTL). These 90 lines were selected among the core collection (164 lines) to maximize the frequency of Col-0 and Cvi-0 at each marker (Table S1). Genotypes were obtained from the Versailles Arabidopsis Stock Center (https://publiclines.versailles.inrae.fr). All lines were grown in two consecutive experiments in the Phenoscope (https://phenoscope.versailles.inrae.fr/), a robotic platform that continually rotates individual pots across a controlled environment chamber, watering each pot according to their weight (Tisné et al., 2013). Two replicates of each RIL were grown in each experiment. For this, seeds were stratified in a 0.1% agar solution for 3 days at 4°C in the dark, and sown on 4 cm plugs of peatmoss substrate mix (blond peat, perlite and sand) wrapped in a non-woven film. Plugs were maintained at saturation (100% of the maximum soil water content) in plastic trays until germination, and immediately after seedlings were thinned on each plug to retain only one individual, with as much synchronous germination as possible across plugs. These plugs were then installed on the Phenoscope robot 8 days after sowing. From sowing to the end of the experiment, plugs were watered twice a day to maintain the soil at 60% of the maximum water content using nutrient solution containing 5mM NO_3_^-^, providing a non-limiting water and nutrient condition.

The photoperiod in the growth room was set to short days (8 h light/16 h dark) to avoid interaction with bolting and to optimize the study of vegetative shoot growth during the exponential growth phase. Light was provided by 6500K white LED sources to an intensity of 230 μmol m^-2^ sec^-1^. The air temperature was set to 21°C during the day and 17°C at night, with a constant relative humidity of 65%. The complete aerial part of each plant was harvested after 23 days (i.e. 31 days after sowing), flash frozen in liquid nitrogen, and grinded for metabolomics. A total of 360 samples were collected for metabolomics.

For QTL confirmation, descendants from 28 heterogeneous inbred lines (HIFs) were grown in the Phenoscope platform. From each HIF we grew three replicates from two independent lines carrying Col-0 alleles at the segregating region and two lines carrying Cvi-0 alleles at the segregating region. Additional replicates were grown for three HIFs for which we only had one Col-0 or Cvi-0 line. The conditions in the Phenoscope were identical to the experiment for the QTL analysis. After 23 days on the robot, the complete aerial part of each plant was harvested, flash frozen in liquid nitrogen, and grinded for metabolomics.

Moroccan accessions Ifr-0 and Khe-0 are originally from the population of South Middle Atlas and Taz-0 from North Middle Atlas; and were kindly provided by Carlos Alonso-Blanco. Accessions F13_6, F13_7, F14_1 and F15_1 originally from the Inferno population in the Island of Fogo were kindly provided by Angela Hancock.

### Metabolite profiling by GC-MS

Plant material was collected in 2 ml Safelock Eppendorf tubes, weighted, flash frozen in liquid nitrogen and ground using the Mixer Mill MM 400 (Retsch®). Fifty mg of powder was transferred into 2 ml Safelock Eppendorf tubes and resuspended in 1 ml of frozen (−20°C) Water:Acetonitrile:Isopropanol (2:3:3) containing Ribitol at 4 μg/ml and extracted for 10 min at 4°C with shaking at 1400 rpm in an Eppendorf Thermomixer. Insoluble material was removed by centrifugation at 20000g for 5 min. Twenty μl were collected and dried overnight at 35°C in a Speed-Vac and stored at −80°C or immediately injected. Three blank tubes underwent the same steps as the samples.

Samples were taken out of −80°C, warmed 15 min before opening, and speed-vac dried again for 1.5 hours at 35 °C before adding 10 μl of 20 mg/ml methoxyamine in pyridine to the samples. The reaction was performed for 90 min at 28°C under continuous shaking in an Eppendorf thermomixer. Fifty μl of N-methyl-N-trimethylsilyl-trifluoroacetamide (MSTFA) (Aldrich 394866-10x1ml) were then added and the reaction continued for 30 min at 37°C. After cooling, 45 μl were transferred to an Agilent vial for injection.

Four hours after derivatization 1 μl of sample was injected in splitless mode on an Agilent 7890A gas chromatograph coupled to an Agilent 5977B mass spectrometer. The column was an Rxi-5SilMS from Restek (30 m with 10 m integraguard column). The liner (Restek # 20994) was changed after each series of 24 samples. Oven temperature ramp was 70°C for 7 min then 10°C/min to 330°C for 5 min (run length 38 min). Helium constant flow was 0.7 mL/min. Temperatures were the following: injector: 250°C, transfer line: 290°C, source: 250°C and quadripole 150°C. Five scans per second were acquired spanning a 50 to 600 Da range. The instrument was tuned with PFTBA with the 69 m/z and 219 m/z of equal intensities. Samples were randomized. Four different quality controls were injected at the beginning and end of the analysis for monitoring of the derivatization stability. An alkane mix (C10, C12, C15, C19, C22, C28, C32, C36) was injected in the middle of the queue for external RI calibration. Injection volume was 1 μl. The instrument was an Agilent 7890B gas chromatograph coupled to an Agilent 5977B mass spectrometer. The column was an Rxi-5SilMS from Restek (30 m with 10 m integraguard column). The liner (Restek # 20994) was changed before the analysis. Oven temperature ramp was 70 °C for 7 min then 10 °C/min to 330 °C for 5 min (run length 38 min). Helium constant flow was 0.7 mL/min. Temperatures were the following: injector: 250°C, transfer line: 290°C, source: 250 °C and quadripole 150 °C. Five scans per second were acquired spanning a 50 to 600 Da range. Instrument was tuned with PFTBA with the 69 m/z and 219 m/z of equal intensities. We performed 43 injections for the samples in the QTL analysis and 9 injections for the samples in the QTL confirmation.

Raw Agilent datafiles were converted in NetCDF format and analyzed with AMDIS (http://chemdata.nist.gov/mass-spc/amdis/). A home retention indices/ mass spectra library built from the NIST, Golm (http://gmd.mpimp-golm.mpg.de/) and Fiehn databases (http://fiehnlab.ucdavis.edu/19-projects/153-metabolite-library) and standard compounds were used for metabolites identification. Peak areas were also determined with the Targetlynx software (Waters) after conversion of the NetCDF file in masslynx format. AMDIS, Target Lynx in splitless and split 30 modes were compiled in one single Excel File for comparison. After blank mean subtraction peak areas were normalized to Ribitol and Fresh Weight.

A response coefficient was determined for 4 ng each of a set of 103 metabolites, respectively to the same amount of ribitol. This factor was used to give an estimation of the absolute concentration of the metabolite in what we may call a “one point calibration”. The values obtained for amino acids were checked by a quantification using ion-exchange chromatography coupled to post-column derivatization by ninhydrin on an Amino-Tac JLC500 (Jeol) after concentrating 10 times the extracts. The values obtained by both methods were found to be very similar.

The U2746 metabolite was characterized in leaves from HIFs using first partial purification on Anion exchange chromatography and second acidic hydrolysis to determine its composition. Anion exchange chromatography were performed on Sep-Pak® Plus Light QMA cartridge (Waters). Cartridges were rinsed with methanol, then 5% ammonium hydroxide, then samples in 5% ammonium hydroxide, then washed with 5% ammonium hydroxide, then washed with methanol and finally eluted with 2% Formic acid in water. Fractions were thoroughly dried in a speedVac® SPD 111V vacuum centrifuge at 45°C until pressure reached less than 100 microns.

Acidic hydrolysis was performed by adding 30μl of 0.5 M TFA in water to the dried samples for 1h at 99°C under constant shaking at 1400 rpm in an Eppendorf Thermomixer. Samples were thoroughly dried in a speedVac® SPD 111V vacuum centrifuge at 45°C until pressure reached less than 100 microns. Samples were then processed as described above.

### Metabolite QTL analysis

All metabolomic values were normalized to the fresh weight of the samples and log2 transformed. Metabolomic analysis in the RIL population resulted in abundance values for 112 metabolites. Six metabolites that were absent in more than half of the samples were removed from the analysis. Values from different derivates of the same metabolite were added and the analysis was performed only for the total amount. We surveyed for metabolite artifacts with the average metabolite concentration of the four replicates for each RIL. Missing values were replaced by the average for the RIL population. Correlation between metabolites was calculated with the heatmap2 function in the gplots package in R (Warnes et al., 2020).

We used the lme4 package in R to extract the concentration of each metabolite in each RIL with a mixed effect linear model with injection as a mixed effect factor and genotype as a fixed factor (Bates et al., 2015). We calculated the effect and LOD score of each marker on each metabolite using the extended Haley–Knott method in the rqtl package (Broman et al., 2003). Thresholds of significance for each metabolite were obtained with a permutation test with 1000 iterations using the extended Haley–Knott method and alpha of 0.01. We assigned QTL positions as the marker that passed the significance threshold and presented the highest LOD score in the chromosome.

### Fine mapping

We performed three rounds of fine mapping on the progeny of heterogeneous inbred family 8HV474 in the conditions mentioned above, with help from IJPB’s Plant Observatory technological platforms. We screened 400, 400, and 800 plants in each round with PCR-based indel, CAPS, or Sanger sequencing markers located in the candidate region (Suppl dataset 4).

Plants for fine mapping were grown in 12×12cm pots in a temperature-controlled glass house supplemented with ∼ 100 μmol m^−2^ s^−1^ artificial light to ensure 16-h photoperiods. Leaf tissue was collected in strips of eight 1.2 ml tubes filled with one metal bead, flash frozen in liquid nitrogen, and disrupted in a ball mill for 30 s (Retsch® MM400, 30 Hz). DNA extraction was performed according to the method proposed by (Edwards et al., 1991).

PCRs were performed in a 20 μL volume containing 20-40 ng genomic DNA, 0.25 U Taq polymerase, 1x supplemented buffer, 0.5 μM forward and reverse primers and 15 μM dNTPs. Bands were visualized on 3% or 4% agarose gels, depending on fragment size differences, containing 4x 10^-5^ vol/vol % ethidium bromide. At each round, selected recombinant lines were allowed to self and their progeny was screened for homozygous lines. The progeny of these fixed lines was used for metabolic profiling in Figure 2. Metabolic profiling was performed as described above, but only normalized values of the interesting metabolites were analyzed.

### Physiological characterization in RILs, HIFs and mutants

Plants from the RIL population were imaged daily during the two consecutive experiments for metabolite QTL analysis in the Phenoscope robotic platform. Images were processed automatically to extract projected rosette area and red/green/blue intensity. Growth rate was calculated as the slope of a generalized linear model using three consecutive projected rosette area values. Significant differences between RILs were calculated with a Student’s T-test comparing the RILs separated by their genotype at c3_09748, the closest marker to the U2746 QTL.

T-DNA insertion mutant lines in AT3G26720 were obtained from http://signal.salk.edu/ with reference numbers SALK_035007C (hereby called s035) and SALK_022207C (hereby called s022). Predicted insertion sites for these mutants are presented in Figure 3.

Metabolomic profiling in the T-DNA insertion mutants and the HIFs described in the previous section was performed in plants grown in 12×12cm pots in walk in growth chambers equipped with fluorescent tubes set at 16/8 h photoperiods and air conditioning set to 21/18 °C cycles. Leaf samples were taken two weeks after flowering, unless otherwise indicated, and processed as described above.

For in-vitro analysis, Arabidopsis seeds were sterilized in a 2% Tween-20 solution for 5 minutes, rinsed with water two times and stratified for three days at 4°C in the dark to promote uniform germination. Seeds where then placed in half strength Murashige and Skoog (MS) medium containing 0, 100, 150 or 200mM NaCl, 1% (w/v) plant agar and 1% (w/v) sucrose. Plates were then placed in a controlled environment chamber with 16/8 h light and 21/19 °C temperature cycles. Plates were scanned in a V600 Epson scanner daily from day 3 to day 7 after sowing. Germination rate was determined by counting the number of germinated seeds from the total number of seeds.

For root growth analysis, seeds were sterilized, stratified and sowed in plates containing half strength MS media as described above. Plates were placed vertically in a growth chamber in the same conditions mentioned above. After three days, similar sized plants were either transferred to a MS agar plate containing 150mM NaCl or to another MS agar plate without salt. After 5 days, plates were scanned in a V600 Epson scanner at 400 ppi and roots were measured using ImageJ.

For analysis of stomata conductance under salt stress, plants were grown in an Aralab walk-in chamber set to 16/8 h light and 21/19 °C temperature cycles. After 30 days, plants were watered with 20ml of a 150mM NaCl solution every three days for two weeks. Stomatal conductance was measured every week in fully expanded mature leaves of a total of 6–9 plants using a steady-state leaf porometer (SC-1, Decagon Devices, LabFerrer, Spain).

### Arabidopsis infection assays using Xanthomonas and Botrytis

The virulent *Xanthomonas campestris* pv. *campestris* (*Xcc*) strain 8004Δ*xopAC* (Guy et al., 2013) was grown on MOKA rich medium (Blanvillain et al., 2007) at 28°C in presence of 50 μg/mL rifampicin. Infection assays were performed as described (Meyer et al., 2005) on four-weeks-old *Arabidopsis thaliana* plants grown in short days (8h light) at 22°C (60% relative humidity, 125 μE.m-2.s-1 fluorescent illumination). Fully expanded leaves were wound-inoculated by piercing three times the central vein with a needle dipped in a bacterial suspension at 10^8^ CFU/mL in 1 mM MgCl_2_. Disease development was scored seven days after inoculation visually in each leaf using the following disease index scale: 0, no symptom; 1, chlorosis at the inoculation site; 2, extended chlorosis; 3, necrosis; 4, leaf death.

For infection with *Botrytis cinerea*, Arabidopsis plants were grown in soil in a growth chamber at 22 °C, 70% humidity, under irradiance of 100 μmol m^−2^ s^−1^ with a photoperiod of 8 h light/16h dark during 5 weeks. *B. cinerea* B05.10 strain was grown on potato dextrose agar at 23 °C under continuous light. After 10 days, each strain produced a dense carpet of conidia. Spores were next washed from the surface of the plate using potato dextrose broth medium. The concentration of spores was determined using a Malassez cell and adjusted to a final concentration of 3.10^5^ conidia/mL. Twenty microliter drops of spore suspension were placed on Arabidopsis leaves of 5-week-old plants. Lesion areas were measured by ImageJ.

### Sequence variation in AT3G26720 in worldwide set of accessions

Sequence variation in AT3G26720 was studied using short read sequences for 1607 worldwide Arabidopsis accessions publicly available at SRA (https://www.ncbi.nlm.nih.gov/sra) with accession numbers PRJEB39079 (334 Cape Verde Islands genomes), PRJEB19780 (73 African genomes), PRJNA273563 (1135 worldwide genomes) and PRJNA646494 (65 Iberian Peninsula genomes). Reads were downloaded and aligned to the TAIR10 Col-0 reference using hisat2 v2.1.0 with default parameters except for the --no-spliced-alignment tag (Kim et al., 2015). We discarded reads with mapping quality lower than 5 using samtools v1.7 (Li et al., 2009). We removed duplicated reads using Picardtools (http://broadinstitute.github.io/picard) and we realigned indels using GATK IndelRealigner (McKenna et al., 2010). We called variants in all 1607 alignments simultaneously using GATK UnifiedGenotyper (McKenna et al., 2010).

For marker design and identification of coding polymorphisms between Col-0 and Cvi-0 in the region of the QTL we obtained short read data from Col-0 and Cvi-0 accessions available at SRA (https://www.ncbi.nlm.nih.gov/sra) with reference numbers SRR492239, SRR013327 and SRR013328. We obtained a list of variants between the two accessions using the same mapping, alignment processing and variant calling methods as above. Primers for small indels and CAPS between Col-0 and Cvi were designed based on the TAIR10 sequence using Primer3 (Untergasser et al., 2012).

For allele reconstruction, we annotated the predicted effect in the protein of all variants localized in AT3G26720 using ANNOVAR (Wang et al., 2010). We then manually curated the list for possible errors in large effect mutations or heterozygous sites. Eighty six percent of the heterozygous sites had a minor allele frequency lower than 25% and were supported by less than 5 reads, so they were converted to homozygous. The rest of heterozygous calls (0.01% of all calls) were discarded. Cases in which two consecutive SNPs were incorrectly annotated were fixed by hand.

## Acknowledgements

This work was supported by a grant from the French National research agency ANR (project ‘StressNet’; ANR-15-CE20-0006-01). This work has benefited from the support of IJPB’s Plant Observatory technological platforms, particularly the Phenoscope platform and VASC stock center. The IJPB benefits from the support of Saclay Plant Sciences-SPS (ANR-17-EUR-0007). We want to thank Carlos Alonso Blanco and Angela Hancock for contributing seeds from Africa and Cape Verde Islands. We would also like to thank Mike Ogden for critical reading of the manuscript. LIPME benefits from the support the ‘Laboratoires d’Excellences’ (LABEX) TULIP (ANR-10-LABX-41) and of the ‘Ecole Universitaire de Recherche’ (EUR) TULIP-GS (ANR-18-EURE-0019).

## Authors’ contributions

FJMR performed all abiotic stress experiments and helped write the manuscript, DW performed the mQTL analysis and contributed to the manuscript; ZX, EG, AR and CC helped with sample preparation and collection in the mQTL experiment; TS and MR performed fine mapping experiments; JM supervised abiotic stress experiments; LDN, EL and AV performed biotic stress experiments; GC supervised all metabolomics experiments and characterized U2746, OL and JMJG conceived the experiments; and JMJG wrote the manuscript, performed bioinformatic analyses and supervised all experiments.

## References

1001 Genomes Consortium (2016) 1,135 Genomes Reveal the Global Pattern of Polymorphism in Arabidopsis thaliana. Cell 166: 481–491

Abdallah MM-S, Abdelgawad ZA, El-Bassiouny HMS (2016) Alleviation of the adverse effects of salinity stress using trehalose in two rice varieties. South Afr J Bot 103: 275–282

Alonso-Blanco C, Bentsink L, Hanhart CJ, Blankestijn-de Vries H, Koornneef M (2003) Analysis of natural allelic variation at seed dormancy loci of Arabidopsis thaliana. Genetics 164: 711–729

Alonso-Blanco C, Gomez-Mena C, Llorente F, Koornneef M, Salinas J, Martínez-Zapater JM (2005) Genetic and molecular analyses of natural variation indicate CBF2 as a candidate gene for underlying a freezing tolerance quantitative trait locus in Arabidopsis. Plant Physiol 139: 1304–1312

Alseekh S, Aharoni A, Brotman Y, Contrepois K, D’Auria J, Ewald J C Ewald J, Fraser PD, Giavalisco P, Hall RD, et al (2021) Mass spectrometry-based metabolomics: a guide for annotation, quantification and best reporting practices. Nat Methods 18: 747–756

Arteaga N, Savic M, Méndez-Vigo B, Fuster-Pons A, Torres-Pérez R, Oliveros JC, Picó FX, Alonso-Blanco C (2021) MYB transcription factors drive evolutionary innovations in Arabidopsis fruit trichome patterning. Plant Cell 33: 548–565

Balasubramaniam T, Shen G, Esmaeili N, Zhang H (2023) Plants’ Response Mechanisms to Salinity Stress. Plants Basel Switz 12: 2253

Barboza L, Effgen S, Alonso-Blanco C, Kooke R, Keurentjes JJB, Koornneef M, Alcázar R (2013) Arabidopsis semidwarfs evolved from independent mutations in GA20ox1, ortholog to green revolution dwarf alleles in rice and barley. Proc Natl Acad Sci U S A 110: 15818–15823

Bates D, Mächler M, Bolker B, Walker S (2015) Fitting Linear Mixed-Effects Models Using lme4. J Stat Softw 67: 1–48

Bianchi G, Gamba A, Limiroli R, Pozzi N, Elster R, Salamini F, Bartels D (1993) The unusual sugar composition in leaves of the resurrection plant Myrothamnus flabellifolia. Physiol Plant 87: 223–226

Blanvillain S, Meyer D, Boulanger A, Lautier M, Guynet C, Denancé N, Vasse J, Lauber E, Arlat M (2007) Plant Carbohydrate Scavenging through TonB-Dependent Receptors: A Feature Shared by Phytopathogenic and Aquatic Bacteria. PLOS ONE 2: e224

Broman KW, Wu H, Sen S, Churchill GA (2003) R/qtl: QTL mapping in experimental crosses. Bioinforma Oxf Engl 19: 889–890

Brotman Y, Riewe D, Lisec J, Meyer RC, Willmitzer L, Altmann T (2011) Identification of enzymatic and regulatory genes of plant metabolism through QTL analysis in Arabidopsis. J Plant Physiol 168: 1387–1394

Chan YF, Marks ME, Jones FC, Villarreal G, Shapiro MD, Brady SD, Southwick AM, Absher DM, Grimwood J, Schmutz J, et al (2010) Adaptive evolution of pelvic reduction in sticklebacks by recurrent deletion of a Pitx1 enhancer. Science 327: 302–305

Cockram J, Mackay IJ, O’Sullivan DM (2007) The role of double-stranded break repair in the creation of phenotypic diversity at cereal VRN1 loci. Genetics 177: 2535–2539

Conklin PL, Norris SR, Wheeler GL, Williams EH, Smirnoff N, Last RL (1999) Genetic evidence for the role of GDP-mannose in plant ascorbic acid (vitamin C) biosynthesis. Proc Natl Acad Sci U S A 96: 4198–4203

Cruz C, Cardoso P, Santos J, Matos D, Sá C, Figueira E (2023) Application of Plant Growth-Promoting Bacteria from Cape Verde to Increase Maize Tolerance to Salinity. Antioxid Basel Switz 12: 488

Durand S, Ricou A, Simon M, Dehaene N, Budar F, Camilleri C (2021) A restorer-of-fertility-like pentatricopeptide repeat protein promotes cytoplasmic male sterility in Arabidopsis thaliana. Plant J Cell Mol Biol 105: 124–135

Durvasula A, Fulgione A, Gutaker RM, Alacakaptan SI, Flood PJ, Neto C, Tsuchimatsu T, Burbano HA, Picó FX, Alonso-Blanco C, et al (2017) African genomes illuminate the early history and transition to selfing in Arabidopsis thaliana. Proc Natl Acad Sci U S A 114: 5213–5218

Edwards K, Johnstone C, Thompson C (1991) A simple and rapid method for the preparation of plant genomic DNA for PCR analysis. Nucleic Acids Res 19: 1349

Elfarargi AF, Gilbault E, Döring N, Neto C, Fulgione A, Weber APM, Loudet O, Hancock AM (2023) Genomic Basis of Adaptation to a Novel Precipitation Regime. Mol Biol Evol 40: msad031

Fernandez O, Béthencourt L, Quero A, Sangwan RS, Clément C (2010) Trehalose and plant stress responses: friend or foe? Trends Plant Sci 15: 409–417

Fujiyama K, Kira Y, Iizuka M, Kimura Y, Seki T (2001) Identification of putative gene encoded on ORF16 of the 81 kb contig of Arabidopsis thaliana chromosome III as alpha-mannosidase. J Biosci Bioeng 92: 401–404

Fulgione A, Neto C, Elfarargi AF, Tergemina E, Ansari S, Göktay M, Dinis H, Döring N, Flood PJ, Rodriguez-Pacheco S, et al (2022) Parallel reduction in flowering time from de novo mutations enable evolutionary rescue in colonizing lineages. Nat Commun 13: 1461

Garcia-Caparros P, Al-Azzawi MJ, Flowers TJ (2023) Economic Uses of Salt-Tolerant Plants. Plants 12: 2669

Guy E, Lautier M, Chabannes M, Roux B, Lauber E, Arlat M, Noël LD (2013) xopAC-triggered Immunity against Xanthomonas Depends on Arabidopsis Receptor-Like Cytoplasmic Kinase Genes PBL2 and RIPK. PLOS ONE 8: e73469

Hooper CM, Castleden IR, Tanz SK, Aryamanesh N, Millar AH (2017) SUBA4: the interactive data analysis centre for Arabidopsis subcellular protein locations. Nucleic Acids Res 45: D1064–D1074

Jakobson L, Vaahtera L, Tõldsepp K, Nuhkat M, Wang C, Wang Y-S, Hõrak H, Valk E, Pechter P, Sindarovska Y, et al (2016) Natural Variation in Arabidopsis Cvi-0 Accession Reveals an Important Role of MPK12 in Guard Cell CO2 Signaling. PLoS Biol 14: e2000322

Joseph B, Atwell S, Corwin JA, Li B, Kliebenstein DJ (2014) Meta-analysis of metabolome QTLs in Arabidopsis: trying to estimate the network size controlling genetic variation of the metabolome. Front Plant Sci 5: 461

Kesari R, Lasky JR, Villamor JG, Des Marais DL, Chen Y-JC, Liu T-W, Lin W, Juenger TE, Verslues PE (2012) Intron-mediated alternative splicing of Arabidopsis P5CS1 and its association with natural variation in proline and climate adaptation. Proc Natl Acad Sci U S A 109: 9197–9202

Kim D, Langmead B, Salzberg SL (2015) HISAT: a fast spliced aligner with low memory requirements. Nat Methods 12: 357–360

Knoch D, Riewe D, Meyer RC, Boudichevskaia A, Schmidt R, Altmann T (2017) Genetic dissection of metabolite variation in Arabidopsis seeds: evidence for mQTL hotspots and a master regulatory locus of seed metabolism. J Exp Bot 68: 1655–1667

Konečná V, Bray S, Vlček J, Bohutínská M, Požárová D, Choudhury RR, Bollmann-Giolai A, Flis P, Salt DE, Parisod C, et al (2021) Parallel adaptation in autopolyploid Arabidopsis arenosa is dominated by repeated recruitment of shared alleles. Nat Commun 12: 4979

Kreiner JM, Giacomini DA, Bemm F, Waithaka B, Regalado J, Lanz C, Hildebrandt J, Sikkema PH, Tranel PJ, Weigel D, et al (2019) Multiple modes of convergent adaptation in the spread of glyphosate-resistant Amaranthus tuberculatus. Proc Natl Acad Sci 116: 21076–21084

Li H, Handsaker B, Wysoker A, Fennell T, Ruan J, Homer N, Marth G, Abecasis G, Durbin R, 1000 Genome Project Data Processing Subgroup (2009) The Sequence Alignment/Map format and SAMtools. Bioinforma Oxf Engl 25: 2078–2079

Lisec J, Meyer RC, Steinfath M, Redestig H, Becher M, Witucka-Wall H, Fiehn O, Törjék O, Selbig J, Altmann T, et al (2008) Identification of metabolic and biomass QTL in Arabidopsis thaliana in a parallel analysis of RIL and IL populations. Plant J Cell Mol Biol 53: 960–972

Loudet O, Gaudon V, Trubuil A, Daniel-Vedele F (2005) Quantitative trait loci controlling root growth and architecture in Arabidopsis thaliana confirmed by heterogeneous inbred family. TAG Theor Appl Genet Theor Angew Genet 110: 742–753

McKenna A, Hanna M, Banks E, Sivachenko A, Cibulskis K, Kernytsky A, Garimella K, Altshuler D, Gabriel S, Daly M, et al (2010) The Genome Analysis Toolkit: a MapReduce framework for analyzing next-generation DNA sequencing data. Genome Res 20: 1297–1303

Meyer D, Lauber E, Roby D, Arlat M, Kroj T (2005) Optimization of pathogenicity assays to study the Arabidopsis thaliana–Xanthomonas campestris pv. campestris pathosystem. Mol Plant Pathol 6: 327–333

Naake T, Zhu F, Alseekh S, Scossa F, Perez de Souza L, Borghi M, Brotman Y, Mori T, Nakabayashi R, Tohge T, et al (2024) Genome-wide association studies identify loci controlling specialized seed metabolites in Arabidopsis. Plant Physiol 194: 1705–1721

Nawaz M, Hassan MU, Chattha MU, Mahmood A, Shah AN, Hashem M, Alamri S, Batool M, Rasheed A, Thabit MA, et al (2022) Trehalose: a promising osmo-protectant against salinity stress-physiological and molecular mechanisms and future prospective. Mol Biol Rep 49: 11255–11271

Neto C, Hancock A (2023) Genetic Architecture of Flowering Time Differs Between Populations With Contrasting Demographic and Selective Histories. Mol Biol Evol 40: msad185

Oliveira DM (2023) Glucuronic acid: not just another brick in the cell wall. New Phytol 238: 8–10

Oliveira DM, Mota TR, Salatta FV, Sinzker RC, Končitíková R, Kopečný D, Simister R, Silva M, Goeminne G, Morreel K, et al (2020) Cell wall remodeling under salt stress: Insights into changes in polysaccharides, feruloylation, lignification, and phenolic metabolism in maize. Plant Cell Environ 43: 2172–2191

Rohman MM, Islam MR, Monsur MB, Amiruzzaman M, Fujita M, Hasanuzzaman M (2019) Trehalose Protects Maize Plants from Salt Stress and Phosphorus Deficiency. Plants Basel Switz 8: 568

Sadak MS (2019) Physiological role of trehalose on enhancing salinity tolerance of wheat plant. Bull Natl Res Cent 43: 53

Saez-Aguayo S, Rondeau-Mouro C, Macquet A, Kronholm I, Ralet M-C, Berger A, Sallé C, Poulain D, Granier F, Botran L, et al (2014) Local evolution of seed flotation in Arabidopsis. PLoS Genet 10: e1004221

Shahid SA, Zaman M, Heng L (2018) Soil Salinity: Historical Perspectives and a World Overview of the Problem. Guidel. Salin. Assess. Mitig. Adapt. Using Nucl. Relat. Tech. Springer International Publishing, Cham, pp 43–53

Shahzad Z, Tournaire-Roux C, Canut M, Adamo M, Roeder J, Verdoucq L, Martinière A, Amtmann A, Santoni V, Grill E, et al (2024) Protein kinase SnRK2.4 is a key regulator of aquaporins and root hydraulics in Arabidopsis. Plant J Cell Mol Biol 117: 264–279

Simon M, Loudet O, Durand S, Bérard A, Brunel D, Sennesal F-X, Durand-Tardif M, Pelletier G, Camilleri C (2008) Quantitative Trait Loci Mapping in Five New Large Recombinant Inbred Line Populations of Arabidopsis thaliana Genotyped With Consensus Single-Nucleotide Polymorphism Markers. Genetics 178: 2253–2264

Tergemina E, Elfarargi AF, Flis P, Fulgione A, Göktay M, Neto C, Scholle M, Flood PJ, Xerri S-A, Zicola J, et al (2022) A two-step adaptive walk rewires nutrient transport in a challenging edaphic environment. Sci Adv 8: eabm9385

Tisné S, Serrand Y, Bach L, Gilbault E, Ben Ameur R, Balasse H, Voisin R, Bouchez D, Durand-Tardif M, Guerche P, et al (2013) Phenoscope: an automated large-scale phenotyping platform offering high spatial homogeneity. Plant J Cell Mol Biol 74: 534–544

Untergasser A, Cutcutache I, Koressaar T, Ye J, Faircloth BC, Remm M, Rozen SG (2012) Primer3--new capabilities and interfaces. Nucleic Acids Res 40: e115

Wang K, Li M, Hakonarson H (2010) ANNOVAR: functional annotation of genetic variants from high-throughput sequencing data. Nucleic Acids Res 38: e164

Wang W, Wang W, Wu Y, Li Q, Zhang G, Shi R, Yang J, Wang Y, Wang W (2020a) The involvement of wheat U-box E3 ubiquitin ligase TaPUB1 in salt stress tolerance. J Integr Plant Biol 62: 631–651

Wang W, Wu Y, Shi R, Sun M, Li Q, Zhang G, Wu J, Wang Y, Wang W (2020b) Overexpression of wheat α-mannosidase gene TaMP impairs salt tolerance in transgenic Brachypodium distachyon. Plant Cell Rep 39: 653–667

Wang YH (2008) How effective is T-DNA insertional mutagenesis in Arabidopsis?

Warnes GR, Bolker B, Bonebakker L, Gentleman R, Huber W, Liaw A, Lumley T, Maechler M, Magnusson A, Moeller S, et al (2020) gplots: Various R Programming Tools for Plotting Data.

Woods R, Schneider D, Winkworth CL, Riley MA, Lenski RE (2006) Tests of parallel molecular evolution in a long-term experiment with Escherichia coli. Proc Natl Acad Sci 103: 9107–9112

Wu S, Alseekh S, Cuadros-Inostroza Á, Fusari CM, Mutwil M, Kooke R, Keurentjes JB, Fernie AR, Willmitzer L, Brotman Y (2016) Combined Use of Genome-Wide Association Data and Correlation Networks Unravels Key Regulators of Primary Metabolism in Arabidopsis thaliana. PLoS Genet 12: e1006363

Wu S, Tohge T, Cuadros-Inostroza Á, Tong H, Tenenboim H, Kooke R, Méret M, Keurentjes JB, Nikoloski Z, Fernie AR, et al (2018) Mapping the Arabidopsis Metabolic Landscape by Untargeted Metabolomics at Different Environmental Conditions. Mol Plant 11: 118–134

Xie KT, Wang G, Thompson AC, Wucherpfennig JI, Reimchen TE, MacColl ADC, Schluter D, Bell MA, Vasquez KM, Kingsley DM (2019) DNA fragility in the parallel evolution of pelvic reduction in stickleback fish. Science 363: 81–84

Yang Y, Guo Y (2018) Elucidating the molecular mechanisms mediating plant salt-stress responses. New Phytol 217: 523–539

Yang Y, Yao Y, Li J, Zhang J, Zhang X, Hu L, Ding D, Bakpa EP, Xie J (2022) Trehalose Alleviated Salt Stress in Tomato by Regulating ROS Metabolism, Photosynthesis, Osmolyte Synthesis, and Trehalose Metabolic Pathways. Front Plant Sci 13: 772948

van Zelm E, Zhang Y, Testerink C (2020) Salt Tolerance Mechanisms of Plants. Annu Rev Plant Biol 71: 403–433

Zhang Z, Ober JA, Kliebenstein DJ (2006) The gene controlling the quantitative trait locus EPITHIOSPECIFIER MODIFIER1 alters glucosinolate hydrolysis and insect resistance in Arabidopsis. Plant Cell 18: 1524–1536

Zhao F, Filker S (2018) Characterization of protistan plankton diversity in ancient salt evaporation ponds located in a volcanic crater on the island Sal, Cape Verde. Extrem Life Extreme Cond 22: 943–954

Zhou H, Shi H, Yang Y, Feng X, Chen X, Xiao F, Lin H, Guo Y (2024) Insights into plant salt stress signaling and tolerance. J Genet Genomics Yi Chuan Xue Bao 51: 16–34

