## Supplemental Figures for "Parallel Evolution of Salinity Tolerance in *Arabidopsis thaliana* Accessions from Cape Verde Islands"

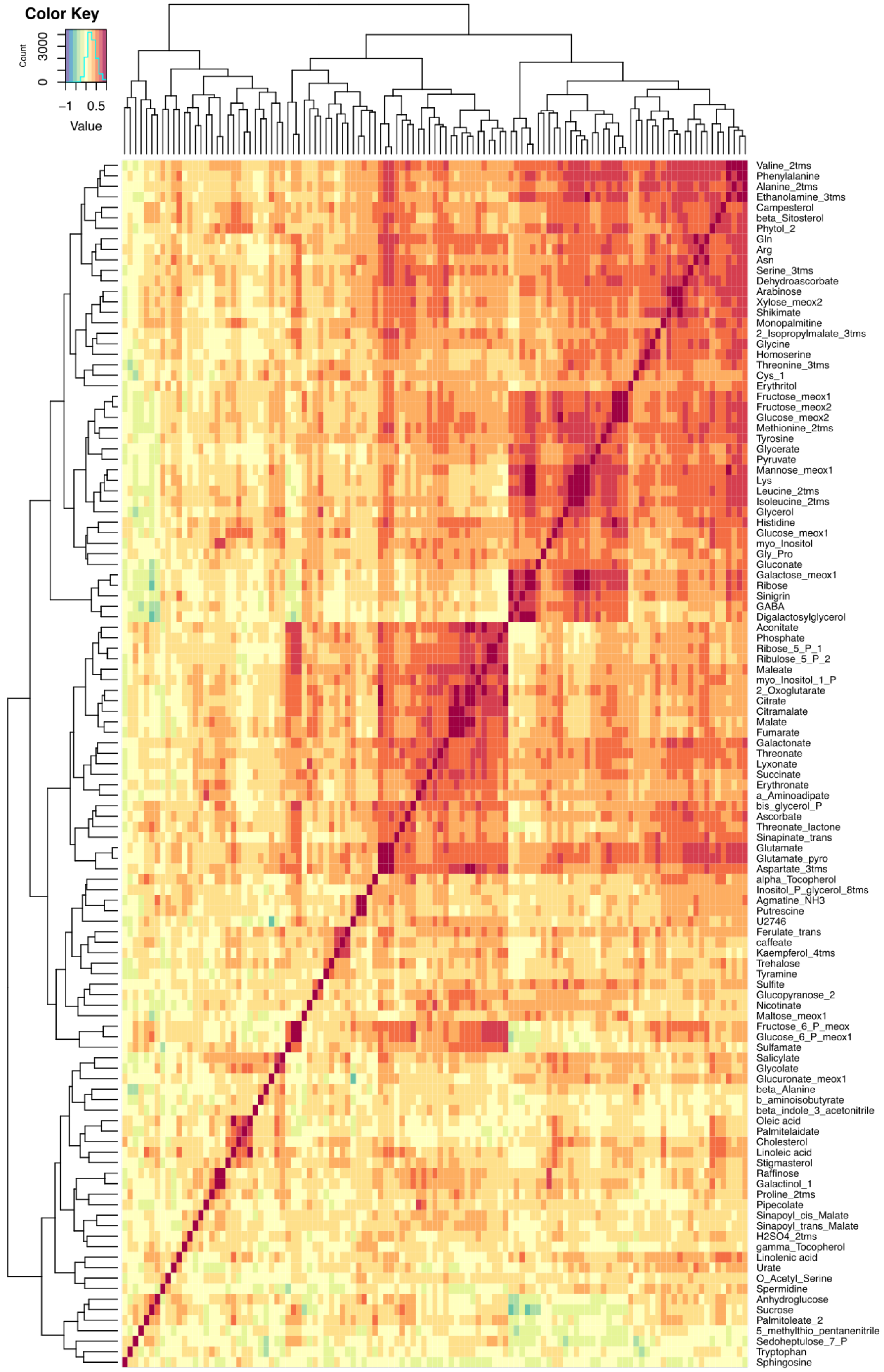


**Figure S1. Metabolite correlation across the RIL population**

Correlation heatmap with all metabolites detected in the RIL population. Average metabolite concentration of all replicates for each RIL was used.

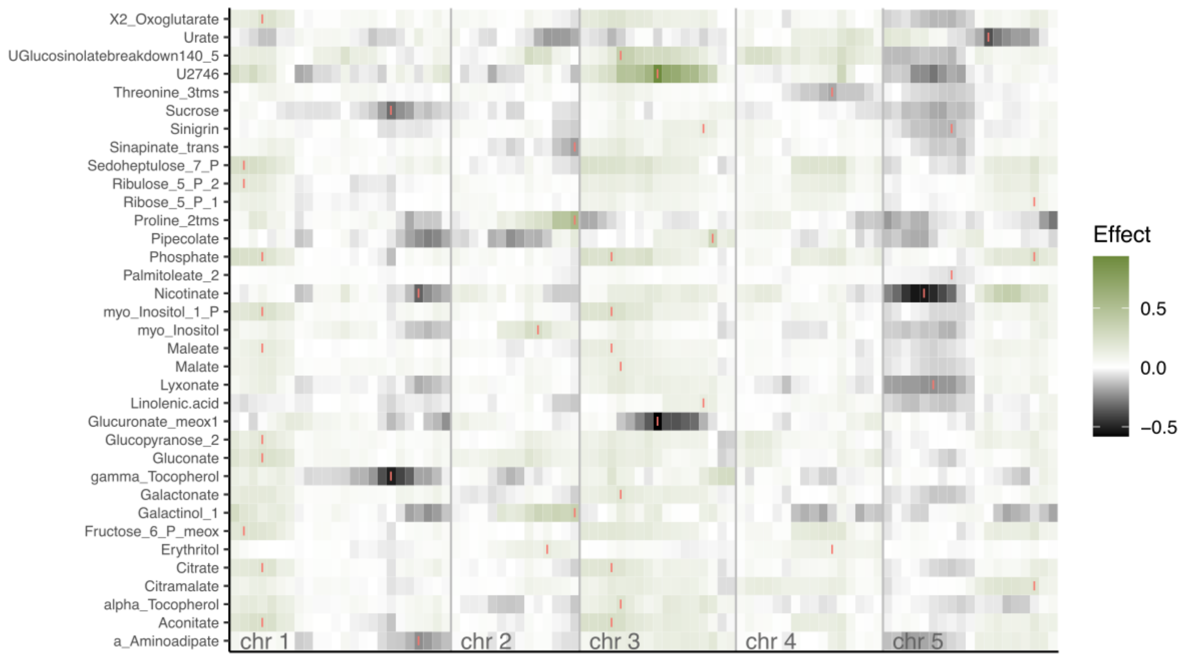


**Figure S2. mQTLs**

Heatmap with QTL effects for all metabolites with at least one QTL with LOD>3. Markers are placed along the x axis and chromosomes are separated by vertical gray bars. Rows represent one metabolite. The color scale in the heatmap represent the additive effects of the homozygous alleles, where positive and negative values indicate higher metabolite abundance caused by the Cvi-0 and Col-0 alleles respectively. QTL positions in the heatmap are indicated with small red vertical lines

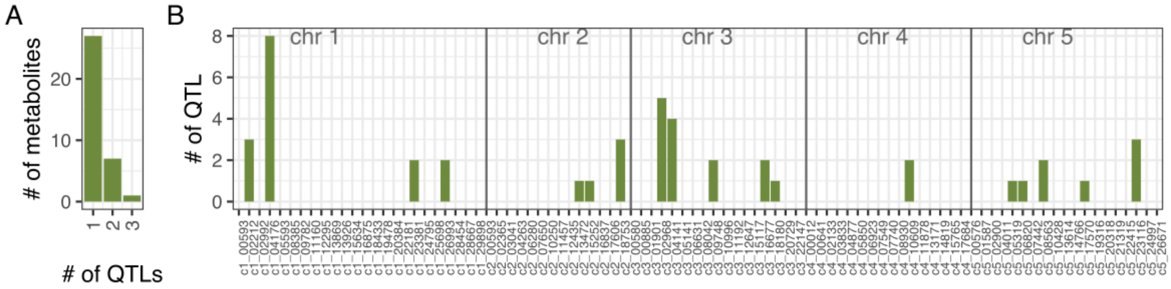


**Figure S3. QTL hotspots**

(A) number of QTLs per metabolite (B) Number of QTLs found at each marker position in the RIL population. Chromosomes are separated by gray vertical bars. Marker names are indicated in the x axis.

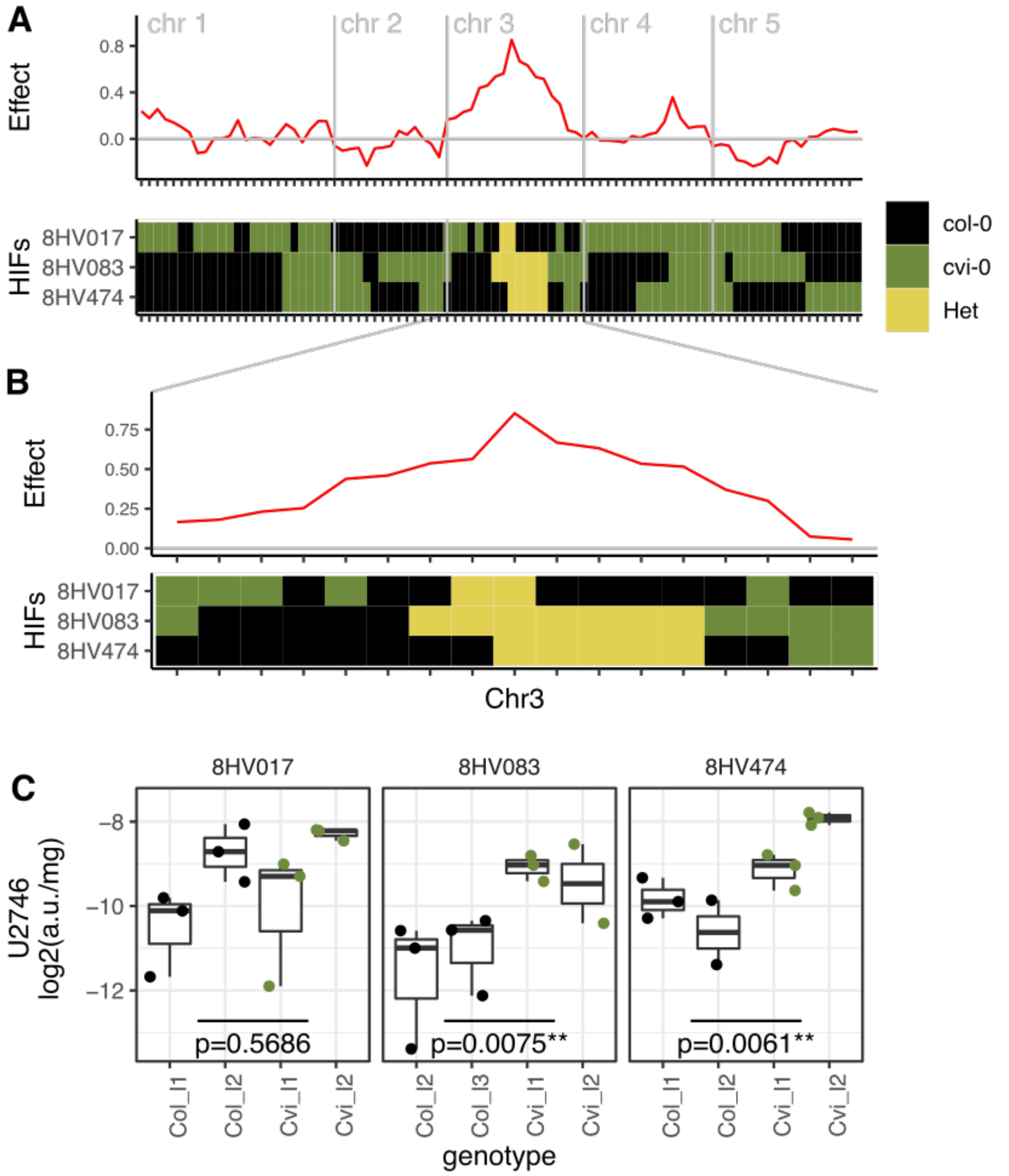


**Figure S4. QTL confirmation using HIFs**

(A) Top: Additive effect of homozygous alleles for U2746. Positive values indicate higher U2746 abundance caused by Cvi-0 alleles. Tick marks in the x axis are the markers used for the QTL analysis and HIF genotyping. Bottom: Genotype for chromosome 3 for three HIFs with a single heterozygous region that overlaps with the U2746 QTL. (B) Zoom on chromosome 3 of the same graphs as above. (C) U2746 abundance in the progeny of each HIF. Y axis represents arbitrary units / mg fresh weight. Plants are homozygous progeny from two independent descendants from each HIF. Numbers in each graph are p values form a one-way ANOVA with genotype as factor. Asterisks mark significant differences ($p < 0.01$).

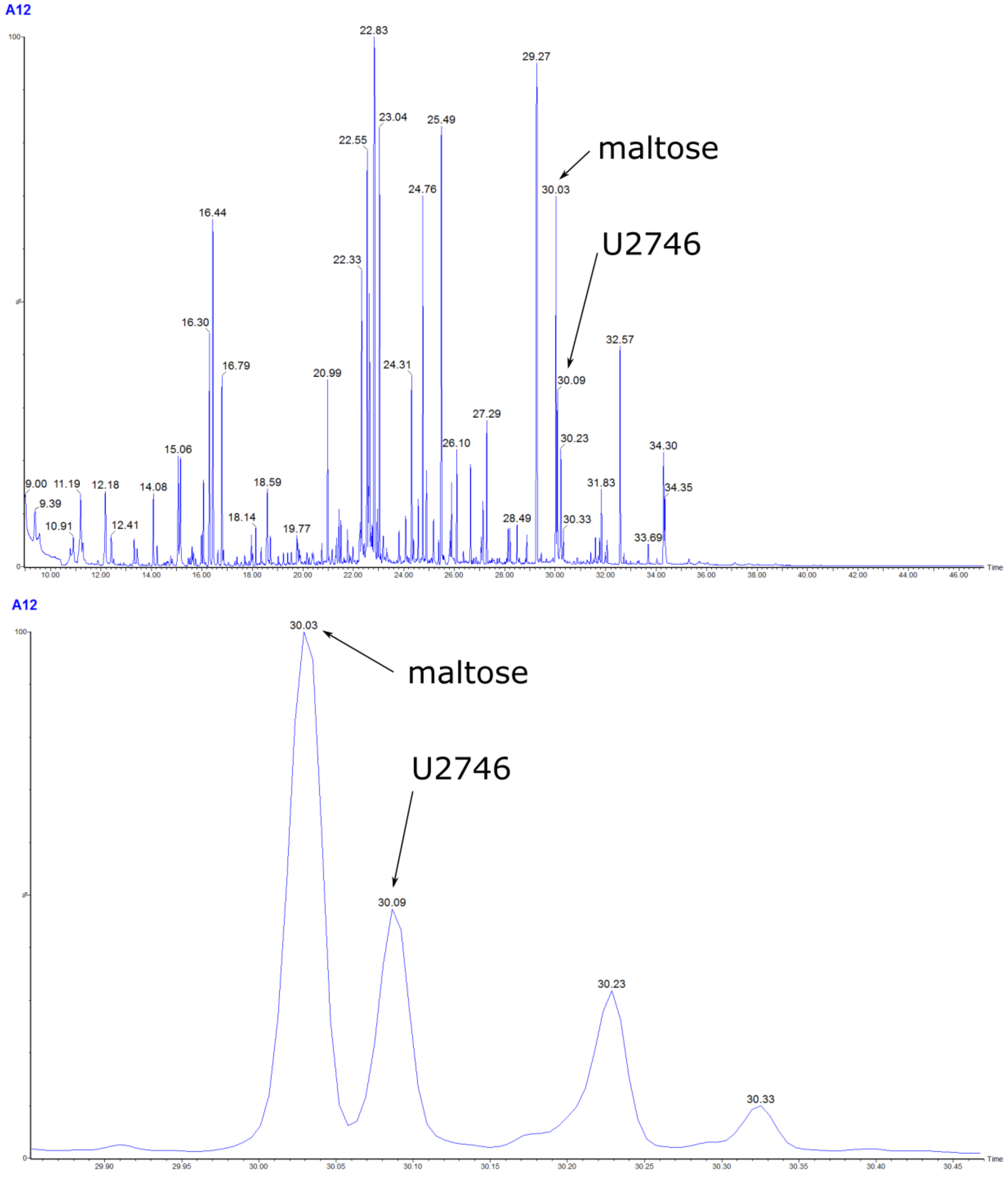


**Figure S5. Retention index U2746**

(A) EI 70ev spectra for U2746 and Maltose from m/z 50 to 1050 from one plant carrying Cvi-0 alleles at the U2746 QTL in chromosome 3. (B) Zoom-in from (A).

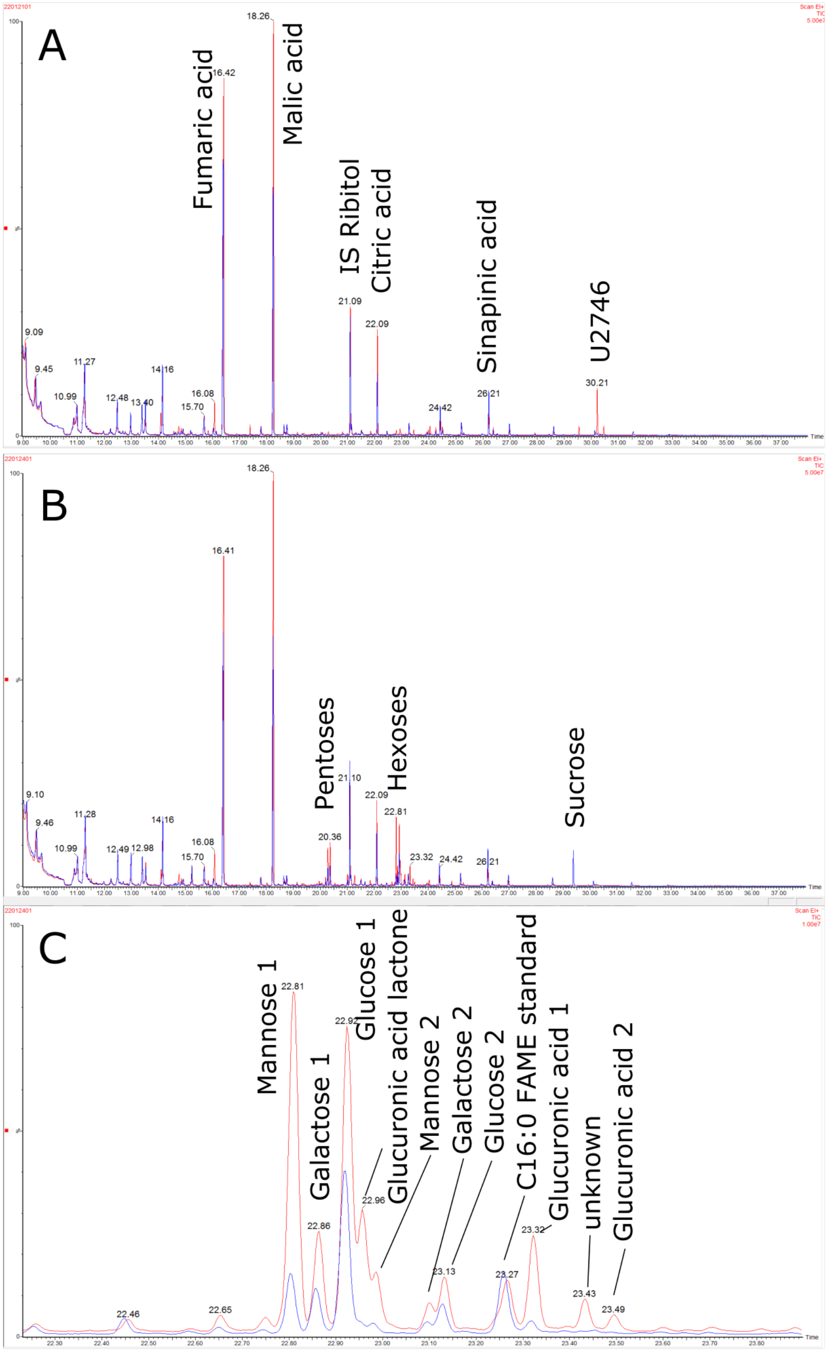


**Figure S6. Fragmentation for U2746**

(A) Purification of the acidic compounds contained in the metabolome extracts of Salk-035 (150mg/ml) and Salk Col-0 (100mg/ml) on a Cation exchange (CEX) Sep-Pak® Plus Light QMA cartridge. (B) 0.5M TFA hydrolysates of the eluates run in 4A.

(B) Ratio of sugars after 0.5M TFA acidic hydrolysis at 60°C, 80°C and 99°C and 1M TFA hydrolysis at 99°C of leaf extracts from HIF-Col and HIF-Cvi.

(C) Zoom of (A) in the 22.4 to 23.9 min region. Known sugar peaks have been marked with the abbreviation of the sugar name followed by 1 or 2 referring to the first and second methoximated isomers. Man, Mannose; Gal, Galactose; Glc, Glucose; GlcA, Glucuronic acid.

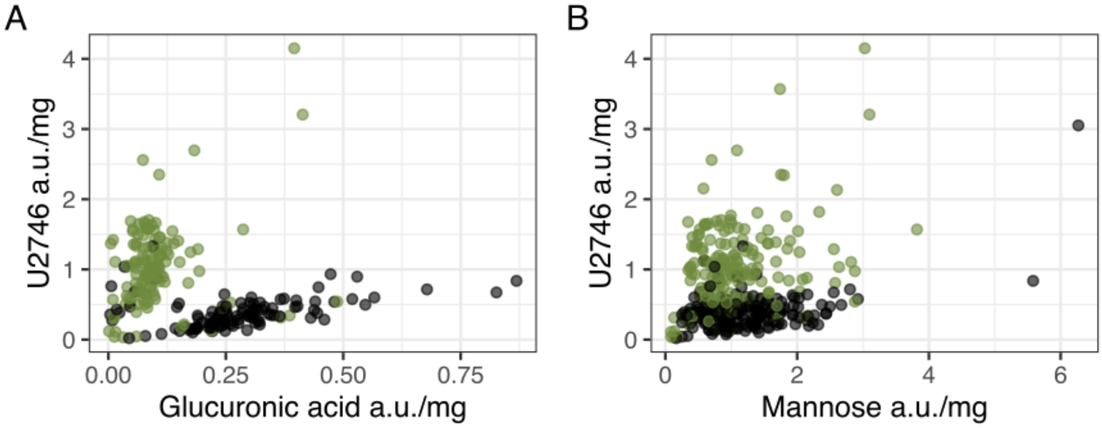


**Figure S7. Correlation between U2746, glucuronic acid and mannose in the RIL population.** Plants in the RIL population were divided by their genotype at marker 09748 (the closest marker to AT3G26720) and the log2 of their normalized concentration for U2746, glucuronic acid and mannose plotted. (A) Comparison between U2746 and glucuronic acid. (B) Comparison between U2746 and mannose.

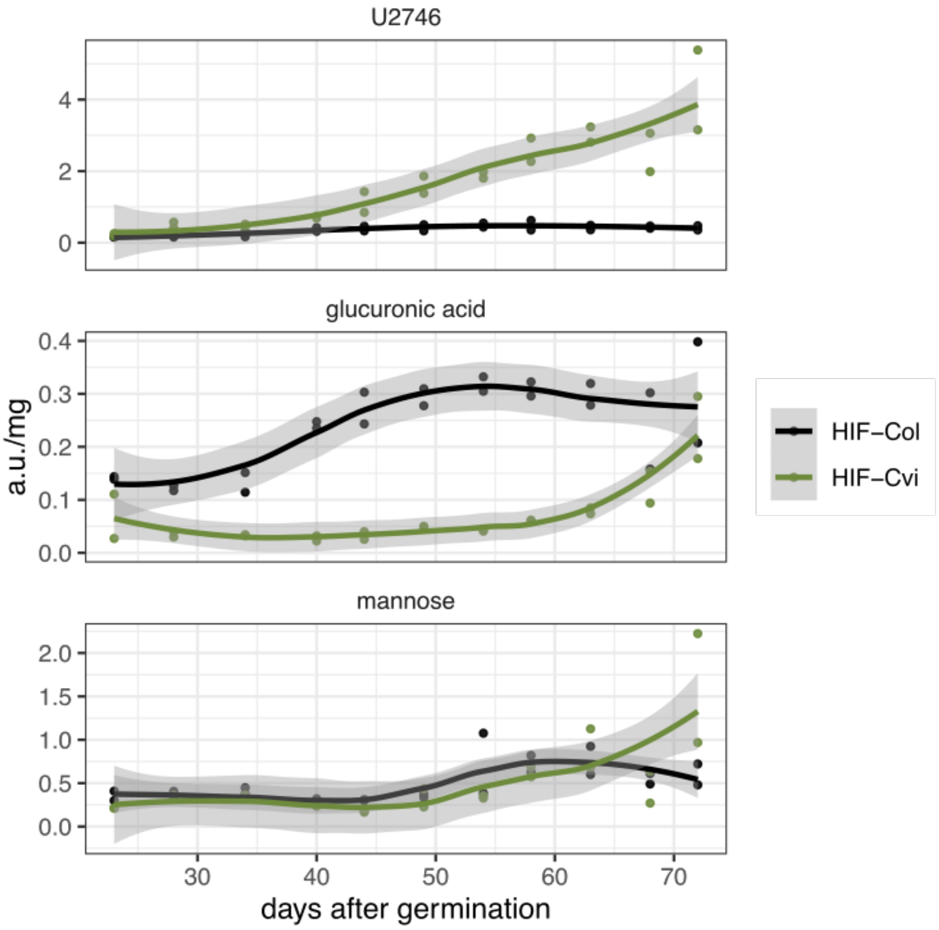


**Figure S8. Developmental time course of U2746**

U2746, glucuronic acid and mannose concentration in HIF-Col and HIF-Cvi from day 23 after planting to senescence. Points represent individual leaf samples in the experiment. Lines are a loess fit per genotype, and the shadowing indicates the 95% confidence level interval for its predictions.

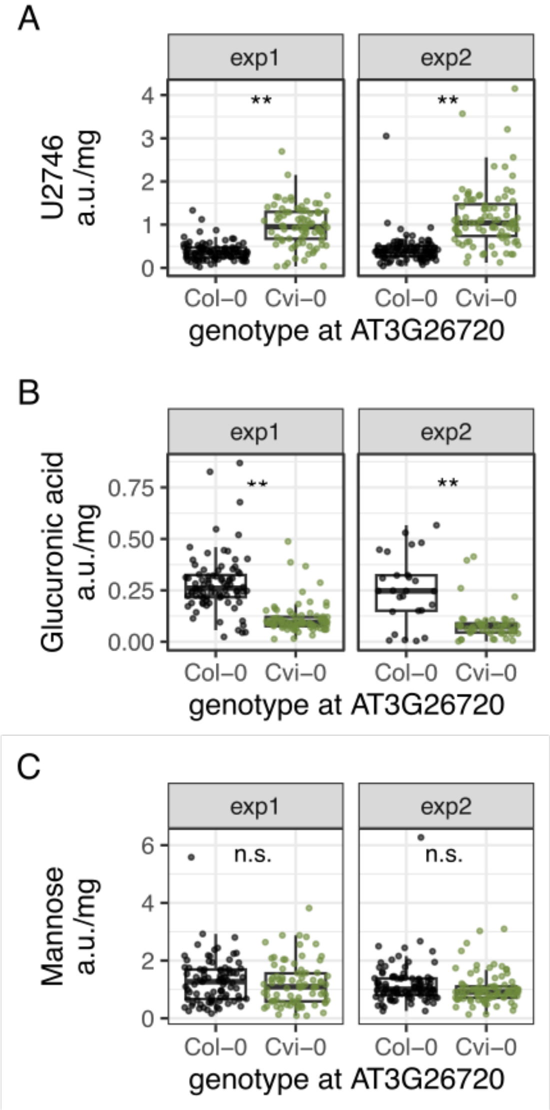


**Figure S9. Metabolite abundance in the RILs grouped by their AT3G26720 allele.** Quantification of U2746 (A), Glucuronic acid (B) and Mannose (C) in samples taken from the RILs during the two experiments for QTL detection in the Phenoscope platform. Two asterisks indicate significant differences between the lines (one way ANOVA, $p<0.01$). n.s. stands for not significant.

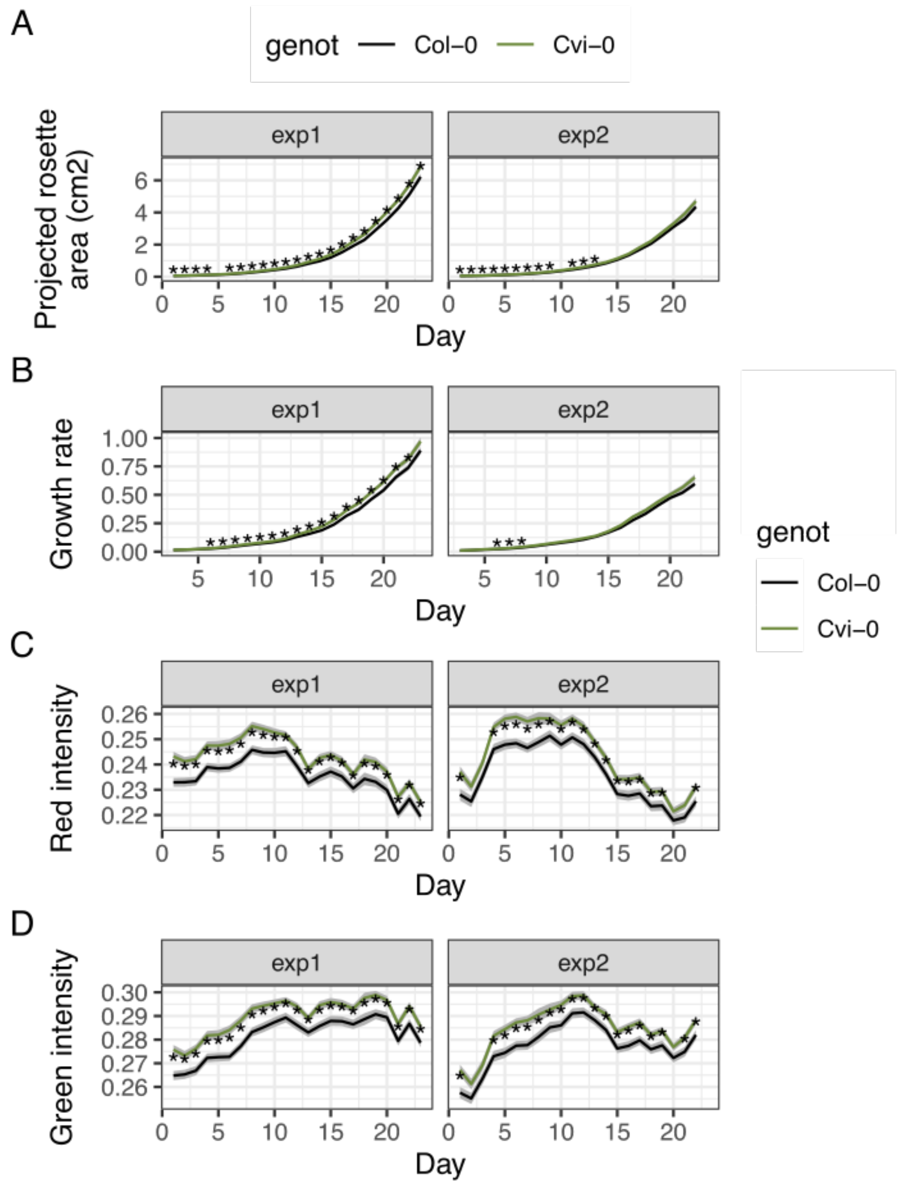


**Figure S10. Characteristics of RILs grouped by their AT326720 allele.**
Quantification of rosette area (A), growth rate (B), red intensity (C) and green intensity (D) from plant images taken from the RILs during the two experiments for QTL detection in the Phenoscope platform. Asterisks indicate significant differences between RILs with Col-0 or Cvi-0 alleles at AT326720 (t-test, $p<0.05$).

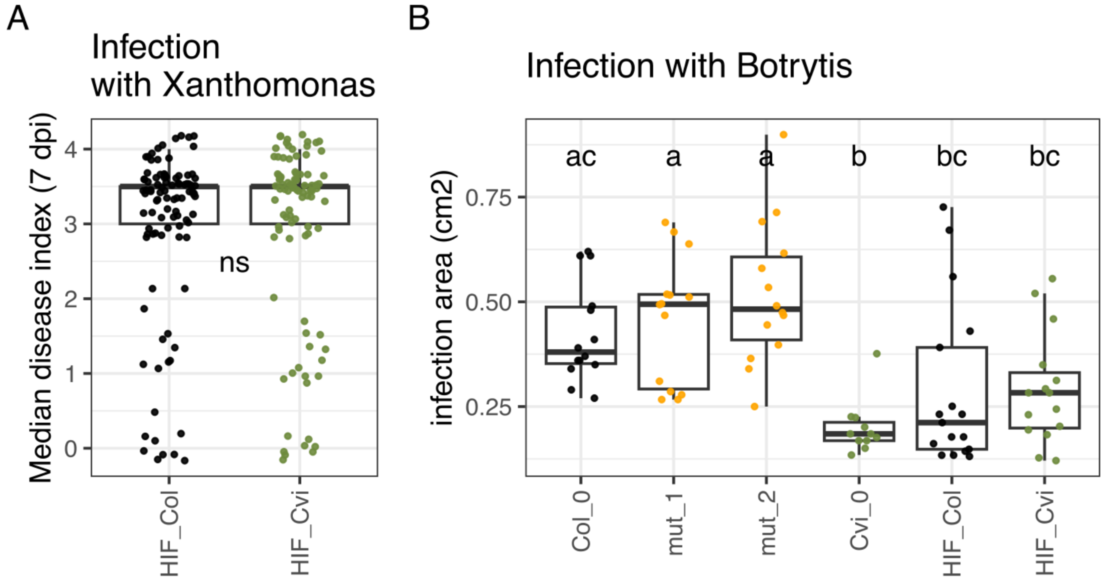


**Figure S11. Effect of AT3G26720 allele in biotic stress responses.**
(A) Virulence assessment taken 7 days post infection in three consecutive experiments. ns stands for not significant differences (Student T-test, p>0.05). (B) Bacterial population density from leaves inoculated with Xanthomonas.

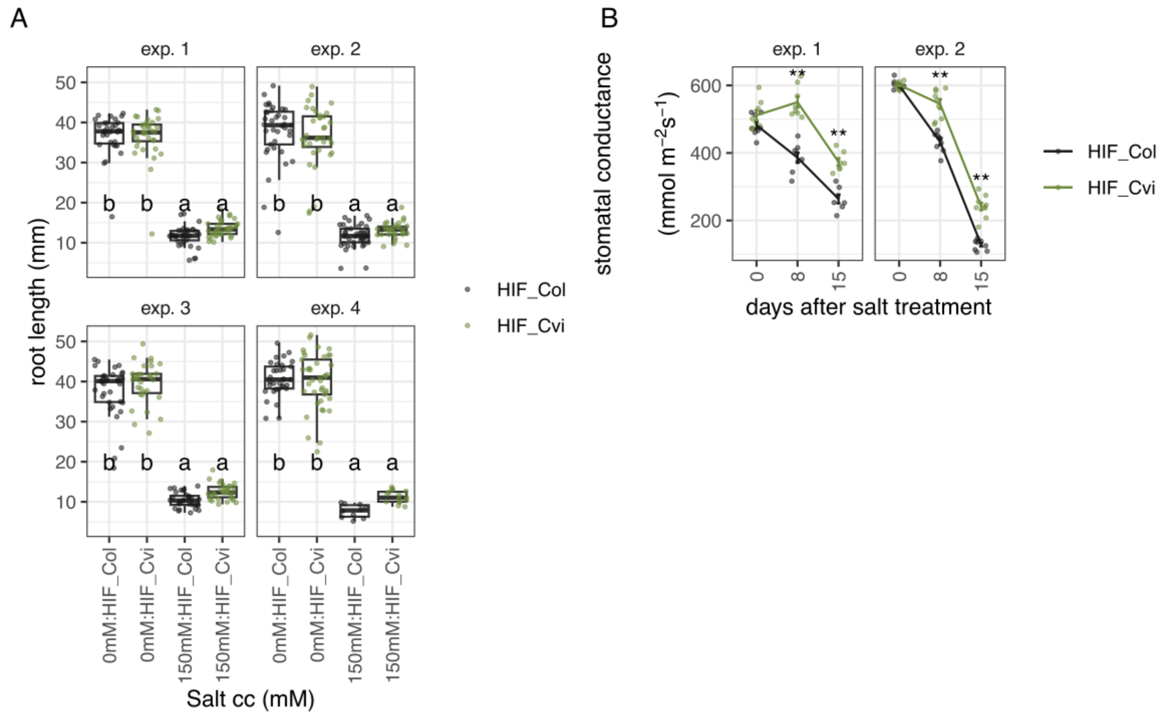


**Figure S12. Effect of mutations in AT3G26720 in responses to high salinity.**
(A) Root length 5 days after transfer to control or salt medium (150mM) in the HIF lines with contrasting alleles at AT3G26720. Data from 4 independent experiments are shown separated. Different letters indicate significant differences between lines and conditions (two-way ANOVA, Tukey HSD test, p<0.05). (B) Stomatal conductance in plants treated with salted water. Treatment starts at day 0. Two independent experiments are shown. Asterisks indicate significant differences at a given day (Student t-test ** p<0.01).

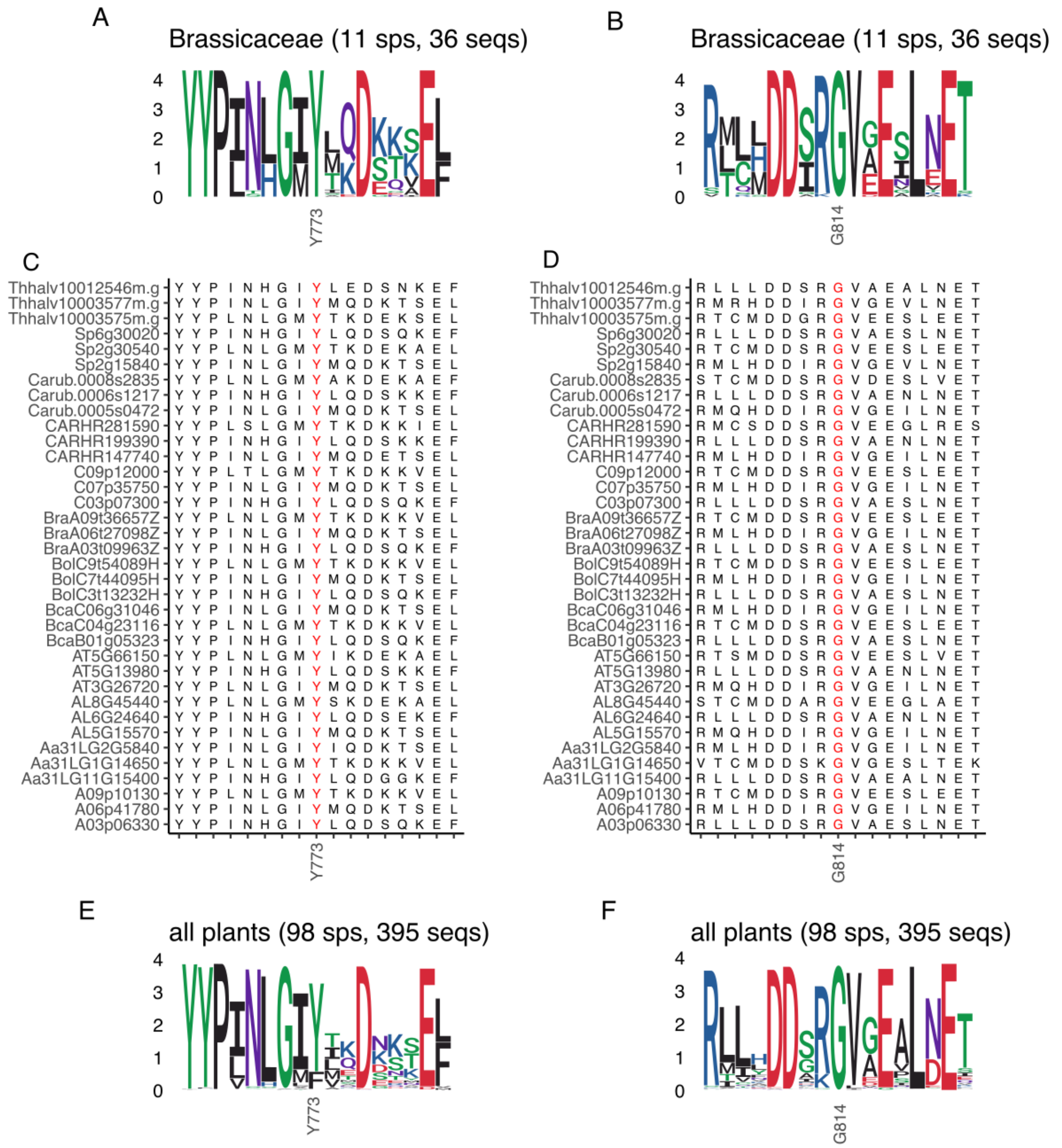

**Figure S13. Allele diversity at positions Y773H and G814D.**
(A) and (B) Sequence logo across 36 orthologous sequences from 11 Brassicaceae species for positions Y773H (A) and G814D (B). (c) and (D) are the corresponding alignments represented in (A) and (B). (E) and (F) Sequence logo across 395 orthologous sequences from 98 plant species for positions Y773H (E) and G814D (F) showing higher conservation for the G814D mutation.

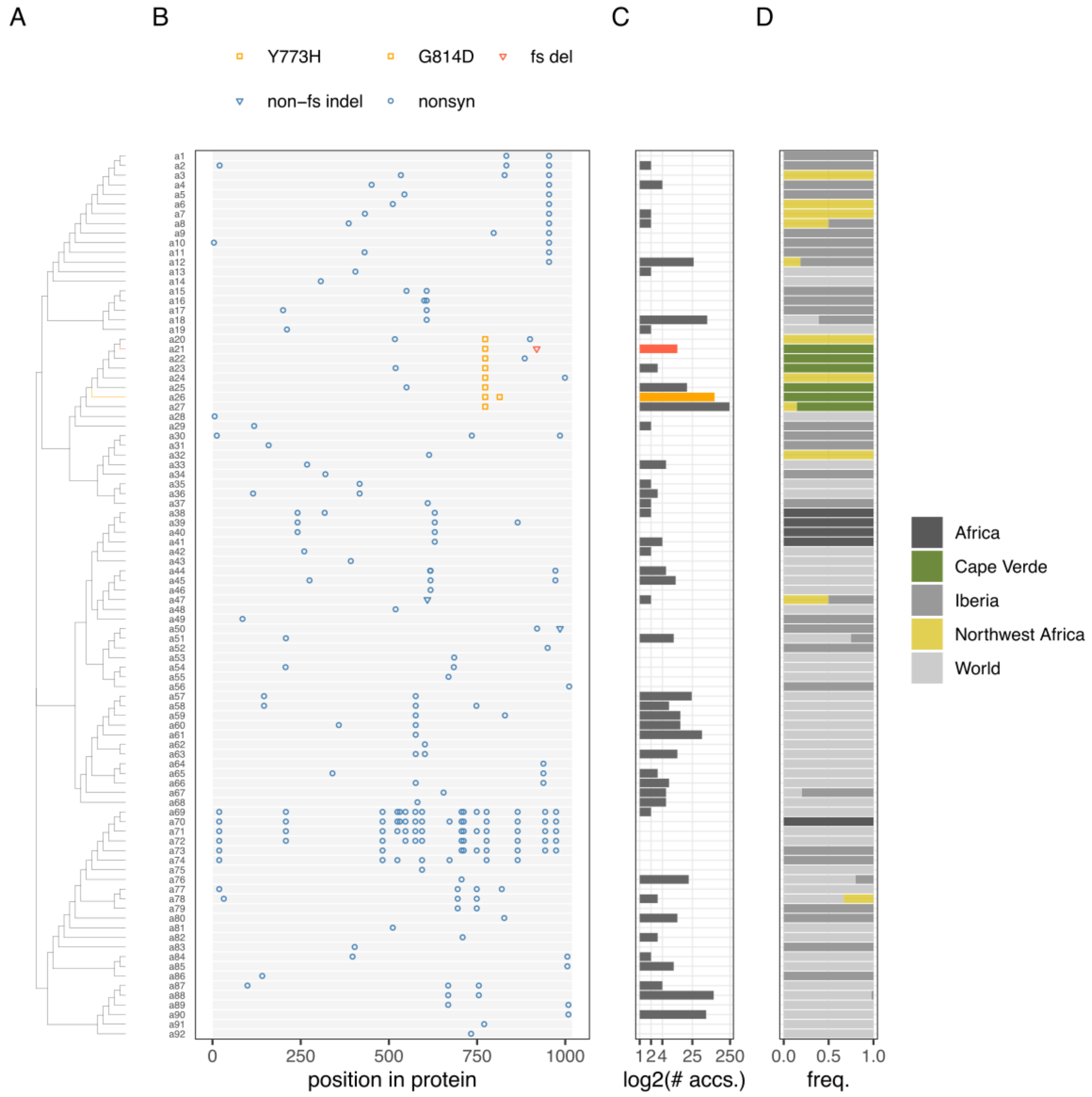


**Figure S14. Allelic diversity at the AT3G26720 protein in Arabidopsis.**

Alleles were reconstructed using homozygous variants from 1607 re-sequenced Arabidopsis accessions. (A) Unrooted maximum likelihood tree from protein sequences. (B) Graphical representation of sequence differences between each allele and the Col-0 reference. In the legend fs stands for frameshift. Blue open circles represent nonsynonymous SNPs. Orange squares represent the Y773H and G824D mutations found in Cvi-0. Red and blue triangles depict frameshift and non-frameshift deletions respectively. (C) Number of accessions carrying each allele in log2 scale. The Cvi-0 allele is colored in orange and the allele carrying a frameshift deletion is colored in red. (D) Frequency of phylogenetic groups for each allele. Both the Cvi-0 allele and the frameshift mutation allele are present only in the Cape Verde Islands.
